# Population persistence under high mutation rate: from evolutionary rescue to lethal mutagenesis

**DOI:** 10.1101/521203

**Authors:** Yoann Anciaux, Amaury Lambert, Ophelie Ronce, Lionel Roques, Guillaume Martin

## Abstract

Populations may genetically adapt to severe stress that would otherwise cause their extirpation. Recent theoretical work, combining stochastic demography with Fisher’s geometric model of adaptation, has shown how evolutionary rescue becomes unlikely beyond some critical intensity of stress. Increasing mutation rates may however allow adaptation to more intense stress, raising concerns about the effectiveness of treatments against pathogens. This previous work assumes that populations are rescued by the rise of a single resistance mutation. However, even in asexual organisms, rescue can also stem from the accumulation of multiple mutations in a single genome. Here, we extend previous work to study the rescue process in an asexual population where the mutation rate is sufficiently high so that such events may be common. We predict both the ultimate extinction probability of the population and the distribution of extinction times. We compare the accuracy of different approximations covering a large range of mutation rates. Moderate increase in mutation rates favors evolutionary rescue. However, larger increase leads to extinction by the accumulation of a large mutation load, a process called lethal mutagenesis. We discuss how these results could help design “evolution-proof” anti-pathogen treatments that even highly mutable strains could not overcome.

## Introduction

Evolutionary rescue (ER) happens when a population confronted with severe stress avoids extinction by genetic adaptation. Understanding and predicting when and how evolutionary rescue occurs is critical in fields as diverse as conservation biology, invasion biology, emergence of new diseases and the management of resistance to treatment in pests and pathogens (see reviews in Gonzalez *et al*. 2013; Carlson *et al*. 2014; Alexander *et al*. 2014; Bell 2017). In all these situations, genetic variation, be it present before the onset of stress, or generated *de novo* after, is a key ingredient for evolutionary rescue, as expected theoretically (e.g. Gomulkiewicz and Holt 1995) and observed experimentally (e.g Ramsayer *et al*. 2013). Because mutation affects both standing and *de novo* genetic variation, it comes as no surprise that a number of evolutionary rescue models, combining stochastic evolution and demography, have predicted that higher mutation rates are associated with higher probability of evolutionary rescue (Orr and Unckless 2008, 2014; Martin *et al*. 2013; Anciaux *et al*. 2018). Few evolutionary rescue experiments have manipulated the mutation rate to test these predictions (reviewed in Bell 2017). For instance, Couce *et al*. (2015) found that two different mutator strains of bacteria with elevated rates of mutations evolved more than 100-fold resistance to antibiotic concentrations that caused the demise of control strains. Mutator alleles are indeed often found in antibiotic resistant strains causing serious health issues (Eliopoulos and Blázquez 2003), raising concern about pathogens escaping our control by evolving higher mutation rates (for theoretical predictions see Taddei *et al*. 1997; Greenspoon and Mideo 2017).

Most mathematical models of evolutionary rescue assume that the population is rescued from extinction by the spread of a single mutant of large effect (Orr and Unckless 2008, 2014; Martin *et al*. 2013; Anciaux *et al*. 2018) and do not describe highly polymorphic populations where several mutations of smaller effects can combine to allow population growth (see however the work of Uecker and Hermisson (2016) and Uecker (2017) where sexual reproduction allows production of such rescue genotypes). The latter situation seems in particular to be common in the evolution of herbicide resistance, especially when the mutational target for resistance is large (Kreiner *et al*. 2018). Even in asexual organisms, when the mutation rate is high, evolutionary rescue may commonly result from the cumulative effect of multiple mutations accumulating stochastically over time in a given lineage. Such a mutation regime is particularly relevant in highly mutable viruses, mutator strains of bacterial (e.g. Springman *et al*. 2010) or cancer cells (e.g. Loeb 2001). Our aim here is to provide theoretical predictions for evolutionary rescue in such a regime with high mutation rates in asexual organisms, complementing existing theory on the subject.

Several complications arise when modelling evolutionary rescue in highly polymorphic populations with high mutation rates. First, the dynamics of allelic frequencies at different loci interact in asexuals. For example, the selective sweep of a given beneficial mutation is hindered by the co-segregation of other beneficial mutations (clonal interference, Gerrish and Lenski 1998). A theoretical study by Wilson *et al*. (2017) recently showed that, when evolutionary rescue is likely, it should most often be driven by soft selective sweeps, where multiple resistance mutations spread through the population simultaneously. Wilson *et al*. (2017) still assumed that each of these lineages carried a single mutation, each with the same effect on the population growth rate. When the mutational target is large, different lineages contributing to rescue are however likely to carry mutations with different fitness effects. Modelling the distribution of mutation effects (as in Martin *et al*. 2013; Anciaux *et al*. 2018) then becomes critical. Finally, when the mutation rate is high, multiple mutations may also accumulate in each lineage, either facilitating evolutionary rescue or impeding it, through their *cumulative* effect. Modelling both beneficial and deleterious mutations and, critically, the epistatic interactions between them, also becomes necessary.

Previous evolutionary rescue theory predicts that higher mutation rate allows populations to withstand higher levels of stress (e.g. Anciaux *et al*. 2018). Yet, there are reasons to expect this prediction not to hold above some critical mutation rate: increased mutation rates also build-up detrimental mutation loads, thus depressing mean fitness despite ongoing adaptation. Indeed, some previous ER models, including both beneficial and deleterious additive effects on growth rates, have found that ER was most likely at intermediate mutation rates (Loverdo and Lloyd-Smith 2013; Greenspoon and Mideo 2017). Artificially increasing the mutation rate has even been proposed as a means to weaken or even eliminate pathogen populations, by a process called lethal mutagenesis (Loeb *et al*. 1999). Models of lethal mutagenesis predict that extinction of the target population could be observed under biologically realistic sets of parameters (Bull *et al*. 2007; Martin and Gandon 2010; Wylie and Shakhnovich 2012). In these models, mean fitness dynamics and extinction stem from the deterministic effects of selection and mutation. Alternatively, Matuszewski *et al*. (2017) discuss the continuity between these models and models of mutational meltdown, where extinction is driven by the interaction of genetic drift and deleterious mutation. Lethal mutagenesis has been investigated empirically for treatment against viruses (Springman *et al*. 2010; Arias *et al*. 2014), bacteria (Bull and Wilke 2008) or cancer cells (Liu *et al*. 2015). In particular, the combination of antiviral treatments with mutagenic agents is investigated as a strategy to fight fast evolving viruses, such as influenza (Bank *et al*. 2016). It seems important to improve our ability to predict whether and when such mutagenic agents will increase treatment efficacy or, conversely, facilitate the evolution of resistance.

The population genetics of adaptation behind the rescue process, in isolated asexual populations, roughly fall into two alternative regimes: rescue may stem (i) from single mutations of large effect (strong selection weak mutation ‘SSWM’ regime) or (ii) from multiple mutations of small effects (weak selection strong mutation ‘WSSM’ regime) (reviewed in Alexander *et al*. 2014). The dichotomy between SSWM and WSSM entails a somewhat simplistic view of adaptation regimes, at the two extremes of all possible mutation rates. The SSWM regime of adaptation has been extensively investigated in population genetics via “origin-fixation” models describing the average behavior of stochastic evolutionary dynamics (McCandlish and Stoltzfus 2014) whereas the WSSM regime has been widely analyzed via deterministic models of quantitative genetics (Lande 1976, 1980). Corresponding evolutionary rescue models further include a coupling of adaptation and demographic dynamics, and naturally fall into the same two regimes (discussed in Anciaux *et al*. 2018). The SSWM regime of evolutionary rescue is characterized by the fact that the first resistant lineage to establish (and thus cause rescue) is only one mutational step from the predominant, sensitive ‘wild-type’ lineage (e.g. Feder *et al*. 2016). Models that describe highly polymorphic dynamics (WSSM regimes, e.g. the quantitative genetic model in Gomulkiewicz and Holt 1995) often use the infinitesimal model assumptions (many unlinked polymorphic loci), which does not apply to asexual populations. In the WSSM regime, the exact stochastic evolutionary dynamics become quickly intractable, and have often been studied by simulation (e.g. Boulding and Hay 2001). Further the latter models often consider initial standing genetic variance as given and pay little attention to the effect of mutation rates in maintaining this variance. They often ignore *de novo* mutations after the onset of stress, on the argument of short timescales being most critical for evolutionary rescue (e.g. Gomulkiewicz *et al*. 2010).

To make analytical progress in our understanding of the effect of mutation rates on the process of evolutionary rescue, we build on two recent theoretical developments (Martin and Roques 2016; Anciaux *et al*. 2018). Anciaux *et al*. (2018) developed a model of evolutionary rescue in the SSWM regime using Fisher’s (1930) geometric model (hereafter “FGM”) to model the distribution of mutation effects on fitness. As in Anciaux *et al*. (2018), we study evolutionary rescue using the FGM. This model assumes a single peak phenotype-fitness landscape, where fitness depends on the position, in phenotype space, of a given genotype relative to an optimum. In the context of ER, stress may affect this landscape in various ways (height, width or position of the peak). In this model, the distribution of mutation effects (both beneficial and deleterious) depend on the context, both genotypic (epistasis) and environmental (e.g. effect of stress). This context-dependence is a key feature of the FGM; it is absent from previous ER models studying the effect of high mutation rates, because they assume additive mutation effects on fitness (e.g. Loverdo and Lloyd-Smith 2013; Greenspoon and Mideo 2017). The variation in the distribution of mutation effects implied by the FGM is qualitatively, and sometimes even quantitatively, consistent with a wealth of empirical observations (reviewed in Tenaillon 2014). Under the assumptions of the FGM, rescue mutants become very rare as the intensity of stress increases, because they require very large mutational steps. As a consequence, Anciaux *et al*. (2018) predict that there is a narrow window of stress levels where the probability of rescue shifts from being very likely to very unlikely. They also predict that this critical level of stress, beyond which adaptation is unlikely, is increased by higher mutation rates (Anciaux *et al*. 2018). Yet, predictions of this model apply to the SSWM regime and may not hold for higher mutation rates.

We extend our previous analysis of evolutionary rescue over Fisher’s geometric model (Anciaux *et al*. 2018) to the more complex and more polymorphic WSSM regime. To do so, we use the approach in Martin and Roques (2016) to model the non-equilibrium dynamics of fitness distributions, in large asexual populations. The form of fitness epistasis assumed may have particular impact on the results because lineages accumulate multiple mutations at different sites over the rescue process. We use the FGM here, which implies a particular form of epistasis that has proven consistent with several observed patterns in fitness epistasis among mutations (Martin *et al*. 2007; Perfeito *et al*. 2014; Blanquart and Bataillon 2016). Moreover, Martin and Roques (2016) showed that, under the FGM, while the fitness dynamics are more complex at higher mutation rates, they are also more predictable and less prone to stochastic fluctuations, even in relatively small populations. To model evolutionary rescue, we still need to describe the demographic stochasticity associated with the extinction process. In the WSSM regime, we thus use a combination of two analytically tractable theories: a deterministic approximation to the dynamics of mean fitness (Martin and Roques 2016) and a diffusion approximation to the stochastic dynamics of population sizes (from Bansaye and Simatos 2015).

Beyond a derivation of the probability of ultimate rescue or extinction, this approach further allows tracking the rescue process over time. As stated in Gomulkiewicz *et al*. (2017), tracking of transient dynamics (population size dynamics, distributions of extinction times) are of high interest for applications of evolutionary rescue theory, yet are not available from existing predictions, which focused mainly on ultimate outcomes. Gomulkiewicz *et al*. (2017) studied the distribution of extinction times for populations doomed to extinction, mostly in the absence of mutation (i.e. with a fixed arbitrary set of competing asexual genotypes at the onset of stress). We extend this analysis to include frequent *de novo* mutation, rescue events involving several mutational steps, a particular form of epistasis and variable mutation effects depending on stress intensity, and an explicit description of the dynamics of mutation load. Our approach captures the continuum from evolutionary rescue to lethal mutagenesis, as mutation rate increases. Interestingly, some parameter ranges prove to greatly limit evolutionary rescue at all mutation rates, i.e. in spite of the possible mutator genotypes.

## Methods

### I. General framework

The present work focuses on an asexual population with independent lineages (e.g. a population of asexual microbes without horizontal gene transfer), facing an abrupt and stressful environmental change (e.g. an antimicrobial treatment). The population initially consists of *N*_0_ individuals with either a single or multiple genotypes and is initially adapted to a non-stressful environment, where its mean growth rate is positive. At the onset of stress, the population is shifted from the non-stressful environment to a stressful environment, where its mean growth rate becomes negative (definition of stress here). In such an environment, in the absence of evolution, the population is doomed to extinction. Evolutionary rescue occurs if at least one resistant lineage (with a positive growth rate in the new environment) establishes, in spite of demographic stochasticity. These resistant mutant lineages can either already be present in the population or arise *de novo* after the onset of stress. It is thus crucial to determine how the number and growth rates of such mutants depend on the new environmental conditions and on the parental genotypes already present in the population. We do so using the FGM detailed below. Note that the main notations used here are summarized in Table 1.

**Table 1:**
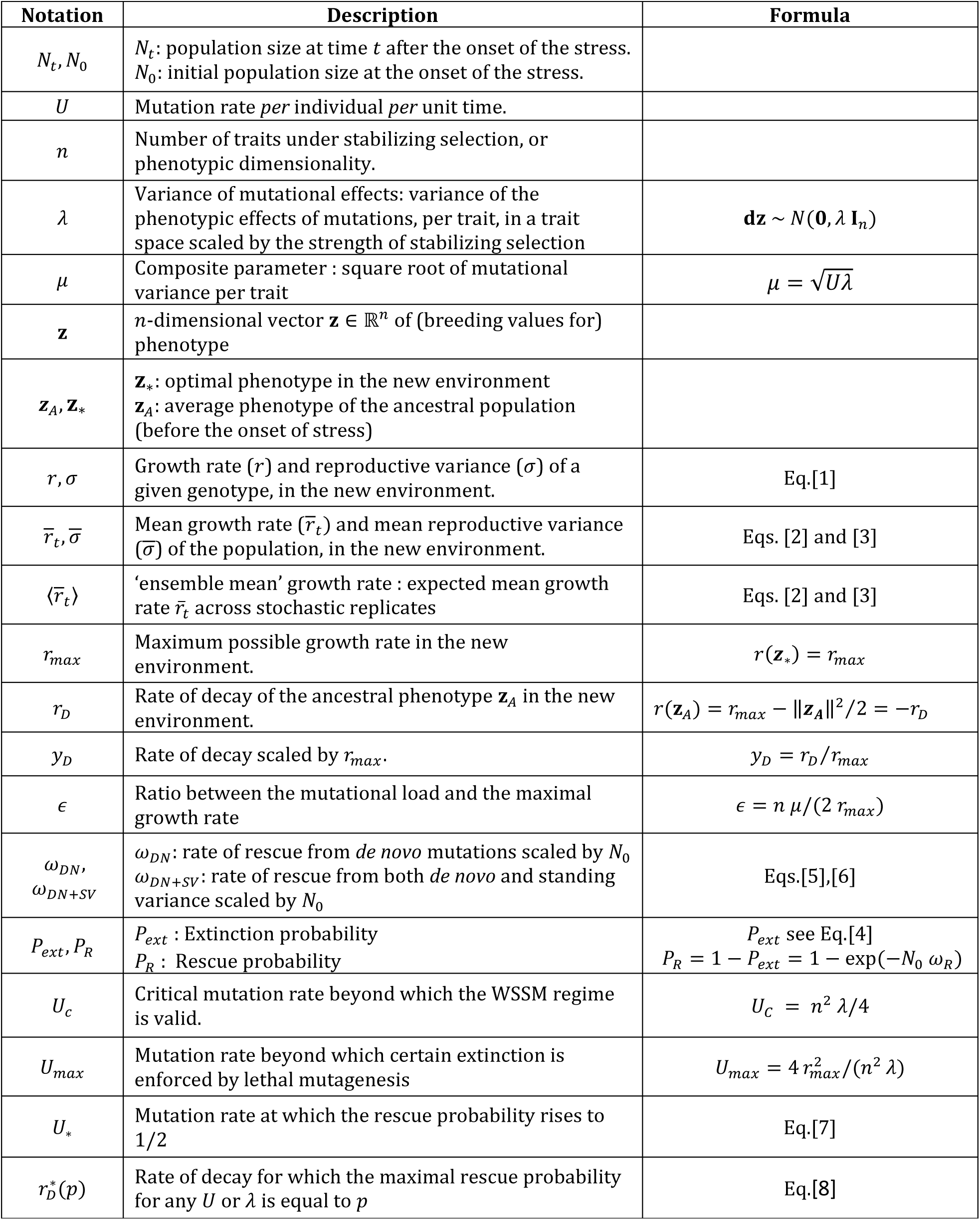
Notations

#### Fitness landscape

In the FGM, a given phenotype is a vector in a phenotypic space of *n* dimensions that determine fitness (here the growth rate *r*). The phenotype of an individual with genotype *i*, is characterized by a vector 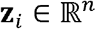 of the breeding values (heritable components) for the *n* traits, and its growth rate is *r_i_*. In a given environment, fitness decays as a quadratic function of the phenotypic distance to a single phenotypic optimum, where the growth rate *r_max_* is maximal at a given absolute level (height of the fitness peak). We assume that each environment is associated with a single optimum and fitness peak. In the scenario investigated here, in the non-stressful environment, the population is close to the ‘ancestral’ optimum **z**_*A*_. When the environment changes, it is assumed to determine a new optimum **z**_*_. Without loss of generality, the height of the peak may also differ between the ancestral and new environments. However, we do require that the *n* dimensions that determine fitness remain the same (in nature and number) across environments. In the new environment, the growth rate of an individual with genotype *i* is given by:

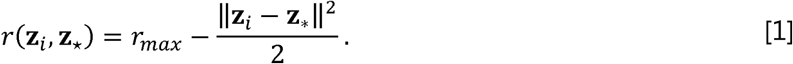

This is an isotropic version of the FGM (all directions are equivalent for selection and mutation) where phenotypes are scaled by selective strength.

#### Stochastic demographic dynamics

We confine our analysis to finite haploid asexual populations. Individuals have independent evolutionary and demographic fate (frequency or density dependence are ignored). Each genotype *i* has a growth rate *r_i_* and a reproductive variance 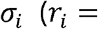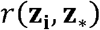, in a given environment with optimum **z**_*_), which define its stochastic demographic parameters in the context of a Feller diffusion approximation (Feller 1951), as in e.g. Martin *et al*. (2013), Gomulkiewicz *et al*. (2017) or Anciaux *et al*. (2018). For simplicity, we further assume here that the average stochastic variance in reproduction is constant over time: 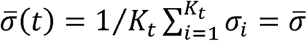 where the average is taken over the *K_t_* genotypes present at time *t*. This can for example be accurate whenever the *σ_i_* are roughly constant across genotypes *i* (discussed in Martin *et al*. 2013 and Anciaux *et al*. 2018).

### II. Evolutionary dynamics

In this section, we describe the model of evolutionary dynamics over the fitness landscape (FGM of the previous section), which is embedded into the ER model. In the following, *de novo* mutations (appearing after the onset of stress) are denoted “DN” and mutations from standing genetic variance (mutants already present before the onset of stress) are denoted “SV”. Correspondingly, evolutionary rescue dynamics from an isogenic population, adapting only from *de novo* mutations, are labelled “DN” and dynamics from a polymorphic population, adapting from both *de novo* mutations and standing genetic variance, are labelled “DN + SV”.

#### 1. Evolutionary dynamics from an isogenic population

The population is maladapted in the new stressful environment and its growth rate is −*r_D_*, corresponding to a decay rate *r_D_*> 0. Mutations arise following a Poisson process with rate *U* per unit time per capita. For a given parent phenotype, each mutation creates a random perturbation ***d*z** on phenotype, which is unbiased and follows an isotropic multivariate Gaussian distribution, 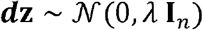 where **I**_*n*_ is the identity matrix in *n* dimensions and *λ* is the variance of mutational effects on traits, standardized by the strength of selection. Mutation effects are additive on phenotype, but not on fitness because *r*(.). is nonlinear (epistasis on fitness and not on phenotype).

In the WSSM regime, the mean growth rate of the population shows limited stochastic variation among replicates, even in reasonably small populations. We thus approximate the evolutionary process by a deterministic fitness trajectory, derived in the WSSM regime under the FGM (Martin and Roques 2016). This seemingly rough approximation can be justified *a priori:* most of the ER process is determined by the speed at which the population adapts at the very onset of stress. This early trajectory takes place when the population is still large and the adaptive process proves to be relatively deterministic, especially over this short timescale, provided that the mutation rate is high enough compared to the mean fitness effect of random mutations (WSSM: 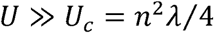, with *n λ*/2 the mean fitness effect of random mutations). Both analytical arguments and simulations detailed in Martin and Roques 2016, showed that mean fitness trajectories are indeed close to the deterministic prediction (with limited variation among replicates), provided that *U* ≫ *U_c_* (WSSM) and *NU* ≫ 1 (large mutational input). Here, we use the deterministic fitness trajectory corresponding to these conditions to approximate the growth rate trajectory of all replicate populations under stress 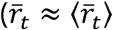, with 〈.〉 the expectation over replicates).

Provided *U* ≫ *U_c_* we thus approximate the trajectory of the mean growth rate of all replicate populations by its deterministic trajectory for the WSSM (Martin and Roques 2016):

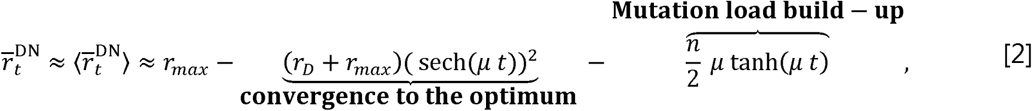

where 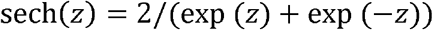 is the hyperbolic secant, 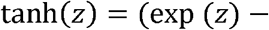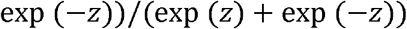 is the hyperbolic tangent and 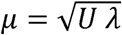 is a composite parameter of the mutational parameters. In Eq.[2] the first term stems from the population nearing the phenotypic optimum, while the second term stems from the build-up of the mutation load as the genetic variance accumulates in the initially clonal population. Recall that *r_D_* is the decay rate of the isogenic population and *r_max_* is the maximum fitness that can be reached in the new environment (with *r_D_* + *r_max_* the fitness distance between the parent genotype’s fitness and the top of the fitness peak). The mean growth rate in Eq.[2] reaches a plateau at infinite time of *r_max_* − *nμ*/2 corresponding to the maximal growth rate minus the mutational load.

#### 2. Evolutionary dynamics from an initially polymorphic population (at mutation-selection balance)

The evolutionary dynamics of rescue from a polymorphic population is obtained by a similar approximation. We assume that the population is initially at mutation-selection balance in the non-stressful environment, with an arbitrary positive mean growth rate. The phenotypic distribution at the onset of stress is centered on a mean phenotype 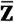, whose growth rate in the new environment is used to characterize the harshness of the stress imposed. For consistency with the isogenic population model above, we thus denote this growth rate 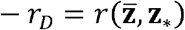, where *r_D_* > 0 is the decay rate of the central genotype as in the previous section. Resistant genotypes may already be present in the population at the onset of stress (“SV”) or appear by *de novo* mutation (“DN”), or arise as combinations of these (multiple step rescue mutants). Using the same reasoning as in the previous subsection, we approximate the mean growth rate of all replicate populations by the deterministic trajectory for the WSSM, i.e. whenever *U* » *U_C_* (Martin and Roques 2016):

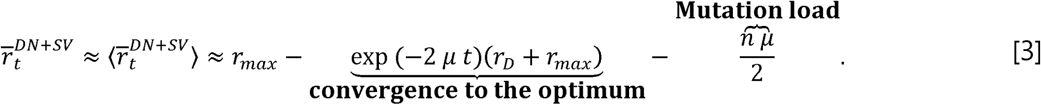

In a polymorphic population at mutation-selection balance, the presence of a mutational load implies that the mean growth rate of the population in Eq.[3] is lower than the mean growth rate of an isogenic population in the same environmental conditions, with the same central genotype 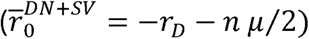. Contrary to Eq.[2], the mutation load is stable (the phenotypic variance is already at equilibrium) while the convergence to the optimum is faster due to the presence of standing variance. The mean growth rate in Eq.[3] ultimately reaches the same plateau as Eq. [2] (the maximal growth rate minus the mutational load).

Note that in both Eqs.[2] and [3] the two mutational parameters *U* and *λ* play entirely symmetric roles through the composite parameter 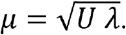 This is because the WSSM regime amounts to a diffusive approximation of the effect of mutation in fitness space (Appendix E.III of Martin and Roques 2016). Like any diffusive approximation, only the mean and variance of the mutation kernel contribute to the dynamics: in this case *Uλ* is the variance induced by mutation during one time unit.

### III. Evolutionary rescue probability

As stated in introduction, we neglect evolutionary stochasticity (i.e. the variance over replicates of the mean fitness dynamic, induced by drift and mutation) but not the demographic stochasticity. The latter is approximated by an inhomogeneous Feller diffusion (see Appendix I section I) with parameters 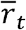 (the expected mean growth rate) and 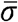 (the reproductive variance averaged across segregating genotypes, which we assume is stable over time): 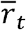 may vary stochastically as genotype frequencies change under the effects of drift, selection and mutation. However, as explained in the previous subsection, we approximate each replicate’s fitness trajectory by its deterministic expectation under the WSSM regime 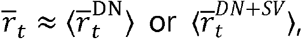 using the relevant cases from Eqs.[2] or [3] (the validity of this approximation is tested against simulations in the Results section). Therefore, the model approximately reduces to a Feller diffusion with constant 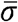 and time-inhomogeneous deterministic growth rate 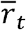. Under these hypotheses, the demographic dynamics follow the stochastic differential equation:

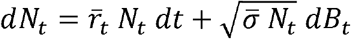 where *B_t_* is a Weiner process(see Appendix I section If or more details). We can then use the results from Theorem 1 of Appendix II (see also Bansaye and Simatos 2015) on inhomogeneous Feller diffusions to derive the probability that the population is extinct before time *t*:

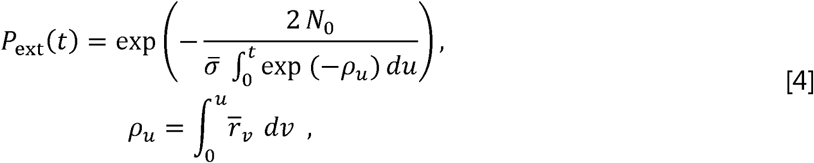

where 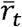 is given by Eqs.[2] or [3]. This expression can be evaluated numerically at any time (*t* > 0. ER happens whenever evolution (change in 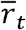) allows extinction to be avoided. Hence, the general form of the rescue probability is readily obtained as the complementary probability of the infinite time limit of Eq.[4], namely the probability of never getting extinct: 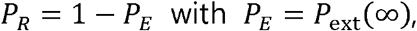 the extinction probability after infinite time. Depending on the scenarios, this rescue probability can be computed explicitly, or approximated via Laplace approximations to the integral 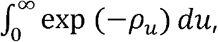 insome parameter ranges (see Results and Appendix I).

### IV. Individual-based stochastic simulations

The analytical predictions are tested against exact stochastic simulations of the population size and genetic composition of populations across discrete, non-overlapping generations (Supplementary Figures 8 shows examples of simulated dynamics of the mean fitness and the population size). Rescue probability was estimated by running 100 replicate simulations until either extinction or rescue occurred. A population was considered rescued when it reached a population size *N_t_* and mean growth rate 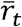 such that its ultimate extinction probability, if it were monomorphic, would lie below 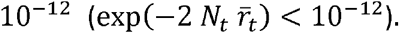 The simulation algorithm is described in Anciaux *et al*. (2018) and also detailed in Appendix I section VIII. Briefly, the number of offspring is Poisson distributed every generation with parameter *e^ri^* for genotype *i*, mutations occur according to a Poisson process with constant rate *U* per capita per generation. The phenotypic effects of the mutations are drawn from a multivariate normal distribution, with multiple mutations having additive effects on phenotype. Fitnesses are computed according to the FGM using Eq.[1]. With such a Poisson offspring distribution, the reproductive variance of genotype *i* is 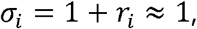 assuming small growth rates *r_i_* « 1, in per-generation time units. So this particular demographic model satisfies our assumption of constant 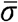 in spite of changes in the genotypic composition of the population: here 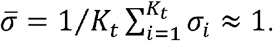 Note also that the analytical derivations (relying on a Feller diffusion (1951) approximation) approximately cover other demographic models (e.g. birth death models, see Martin *et al*. 2013; Gomulkiewicz *et al*. 2017), as long as they also satisfy 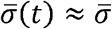 constant.

For *de novo* rescue, the initial population consisted of a single genotype optimal in the environment before stress. When considering contribution from standing variance, the initial population was at mutation selection balance in the former environment (before the onset of stress). More precisely, 10 replicate initial equilibrium populations were generated and 100 replicate simulations were run from each of these populations. The overall rescue probability is then the average, over the 10 equilibrium populations, of the rescue probabilities estimated by the proportion of rescued populations over the 100 simulations.

### V. Appendices and supplementary information

Analytical derivations and supplemental figures are described in Appendix I. Appendix II contains the derivations for the inhomogeneous Feller diffusion. Supplementary file 1 provides the details of the analytical derivations, the code producing the figures and the simulation code as a Mathematica© .cdf file (MATHEMATICA v. 11.3 Wolfram Research 2018) which can be open using the free “CDF player” available on the Wolfram website. Supplementary file 2 provides the Matlab© (MATLAB 2015a, The MathWorks, Natick, 2015) source code for the curve fitting procedure used for Eq.[8].

## Results

### Rescue probability: general form

In spite of their difference in the population genetics underlying ER, the two scenarios with or without standing variance yield similar expressions for the probability of ER (as shown in Eqs.(A5) and (A11-A12) in Appendix I):

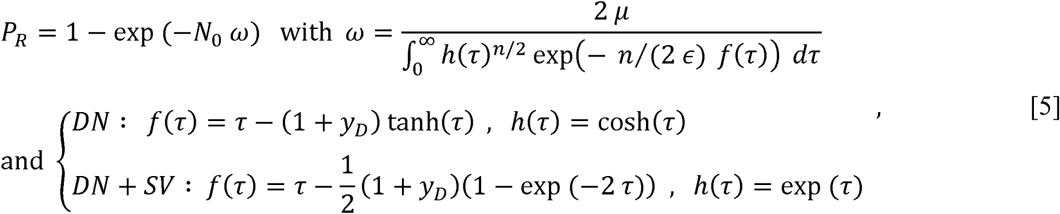

where the particular form of the functions *f*(.). and *h*(.) depend on the chosen scenario. Here, 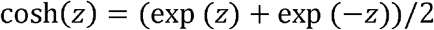 is the hyperbolic cosine and we introduce two scaledparameters: 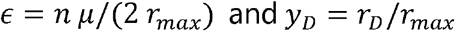 (recall that 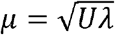 dependsonboth mutation rate and effect). The parameter *y_D_* describes how fast the initial clone decays, compared to how fast the optimal genotype grows, it gives a scaled measure of the harshness of the stress imposed (see also Anciaux *et al*. 2018). The parameter is ∊ the ratio of the mutation load (at mutation-selection balance) and the maximal absolute growth rate that can be reached in the stress. Certain extinction by lethal mutagenesis occurs whenever the load is equal to or larger than the maximal growth rate (i.e. ∊ ≥ 1). A small value of ∊ means that we are far from this certain extinction regime.

The quantity ω in Eq.[5] is akin to a ‘per lineage rate of rescue’. Analytical expressions for this rate are either unavailable (*DN*) or complicated (*DN* + *SV*), Eq.(A12) in Appendix I).

### Weak selection, intermediate mutation approximation

To get a more direct insight into the impact of each parameter, we sought an approximate expression for ω, detailed in Appendix I section III and IV. This approximation applies with an intermediate mutation rate, where ER mainly depends on the early adaptation of the population to stress, and not on the ultimate mutation load (lethal mutagenesis). More precisely, it requires that the WSSM approximation be accurate while ∊ remains small, which implies a small λ and intermediate: 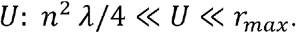 This is why we call this range a ‘weak selection, intermediate mutation’ approximation (WSIM in Figure 1 and 2).

**Figure 1:**
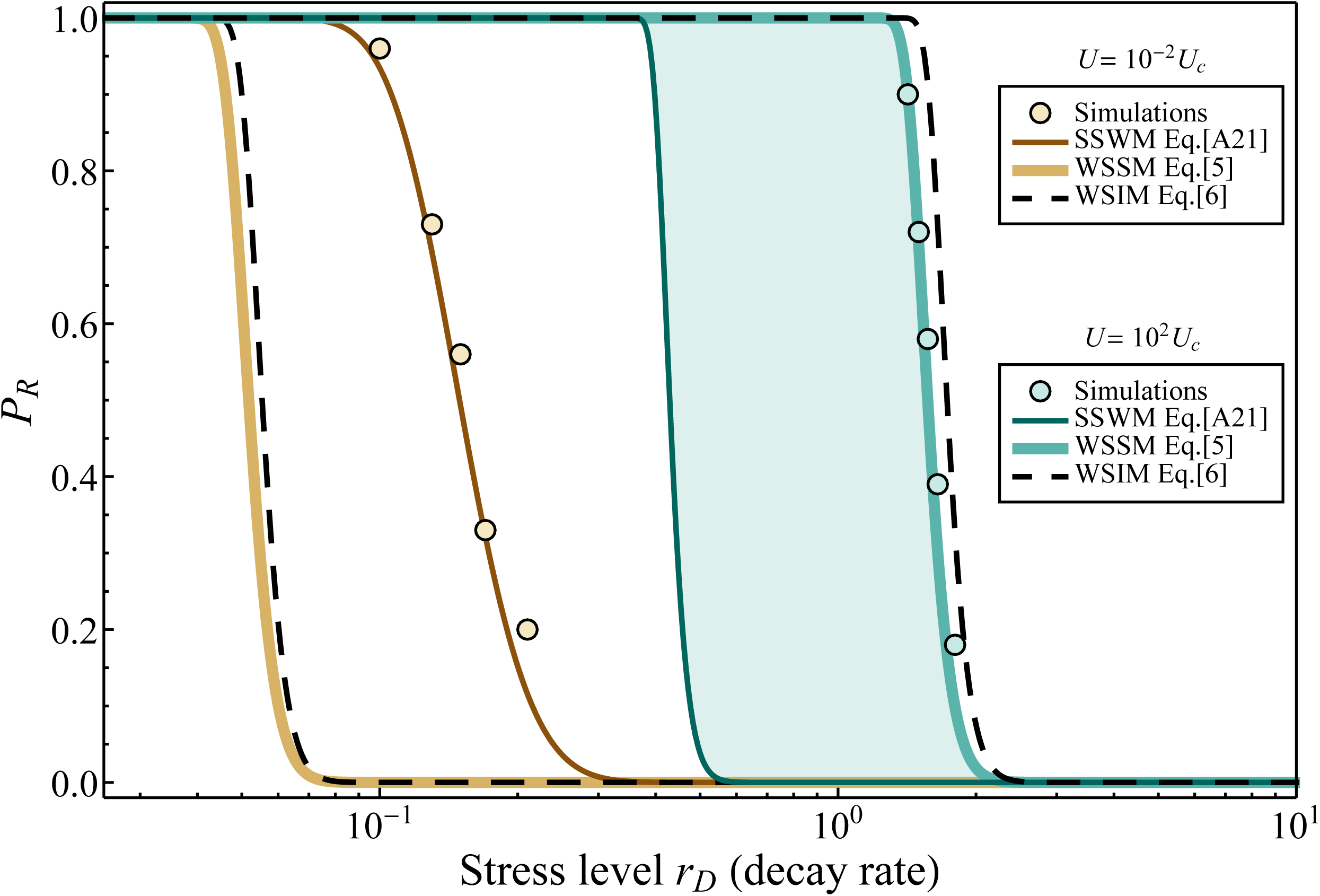
ER probability against decay rate *r_D_* for a population without standing variance (*DN*). Dots show the results of 10^2^ simulations; thin plain lines: Eq.(5) from Anciaux *et al*. 2018 (recalled in Appendix Eq.(A21)) derived under the SSWM regime; thick plain lines: Eq.[5] derived under the WSSM regime; dashed lines: the corresponding closed form expression in Eq.[6] derived under the weak selection intermediate mutation regime (WSIM). The shaded area corresponds to the extra contribution to ER from multiple mutants, compared to single mutants. All models and simulations are shown for a high mutation rate (blue) or a low mutation rate (brown), indicated in legend. As expected, the model from Anciaux *et al*. (2018) derived in the SSWM regime captures the simulations at low mutation rates (*U* = 10^−2^ *U_c_*, brown), whereas Eqs.[5] and [6] captures the simulations at high mutation rates (*U* = 10^2^ *U_c_*, blue). Other parameters are *N*_0_ = 10^5^, *n* = 4, *r_max_* = 1 and *λ* = 5 × 10^−3^.

Under this approximation, the *per capita* rate of rescue ω takes a roughly similar form for both scenarios with or without standing variance (detailed in Eqs.(A8) through (A14) in Appendix I):

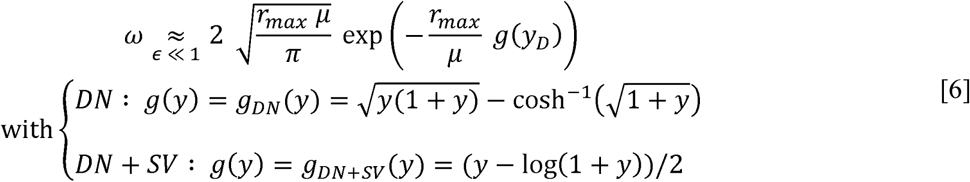

In both cases, the function *g*(.) is positive and increases (roughly log-log linearly) with *y_D_* = *r_D_/r_max_*. The accuracy of this approximation is illustrated in Supplementary Figures 2 and 3 and Figures 1 and 2. Note that, whenever the approximation applies, the ER probability is independent of dimensionality (*n*). This directly stems from the fact that the growth rate trajectories in Eqs.[2] and [3] only depend on dimension via the ‘mutation load’ terms, whose contribution is negligible when far from the lethal mutagenesis threshold (*U* « *r_max_*).

**Figure 2:**
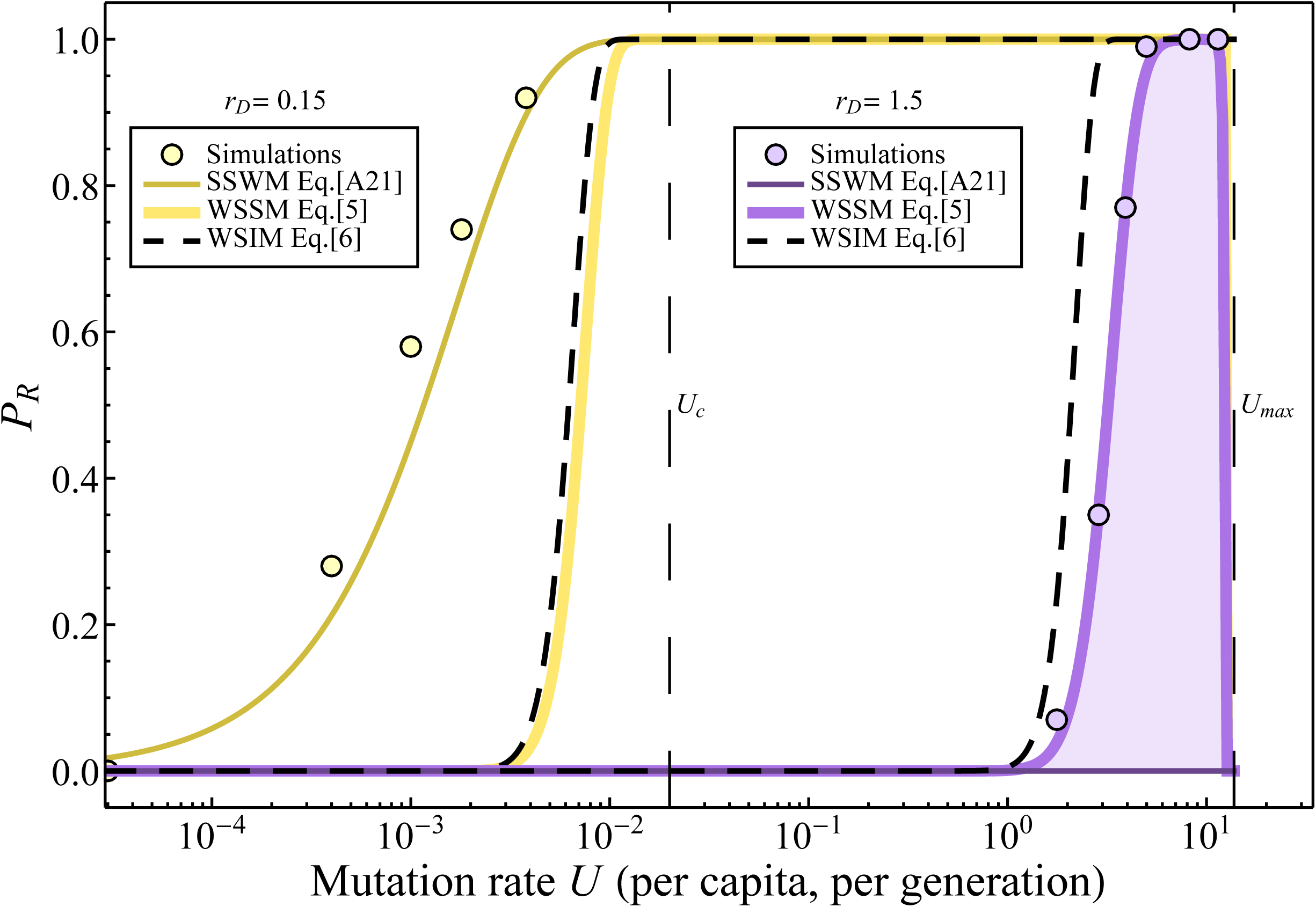
ER probability against mutation rate for a population without standing variance (). Dots show the results of simulations; thin plain lines: Eq.(5) from Anciaux *et al*. 2018 (recalled in Appendix Eq.(A21)) derived under the SSWM regime; thick plain lines: Eq.[5] derived under the WSSM regime; dashed lines: the corresponding closed form expression in Eq.[6] derived under the weak selection intermediate mutation regime (WSIM). The shaded area corresponds to the extra contribution to ER from multiple mutants, compared to single mutants. All models and simulations are shown for a high decay rate (purple) or a low decay rate (yellow), indicated in legend. For low mutation rates, corresponding to the SSWM regime, simulations show that populations handle only low stresses, which is captured by the model from Anciaux *et al*. (2018) derived in the SSWM regime. Whereas for high mutation rates, corresponding to the WSSM regime, simulations show that populations handle high stresses, which is captured by the present model derived in the WSSM regime. Other parameters are,, and

### Sharp decay in ER probability with increasing stress levels

A possible measure of stress intensity in ER is the rate of decay *r_D_* of a population after the environmental change (see also Anciaux *et al*. 2018). However, stress might also affect other parameters of the FGM: the height of the fitness peak *r_max_*, the mutation rate *U* or the variance of mutational effects: *λ*. We detail the two latter effects (which affect ER via the composite parameter 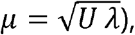, in the next section, and focus here on *r_max_* and *r_D_*. In the following we use the per capita rate of rescue *ω*_*DN*_ from Eq.(5) of Anciaux *et al*. (2018) in the SSWM regime (recalled in Appendix I section VIII, Eqs.(A20)-(A21)), as a comparison to the present results in the WSSM regime. By basic properties of the FGM described in Anciaux *et al*. (2018), increased *r_D_* means both a faster decay (purely demographic effect of stress) and a larger shift in optimum (which affects the whole distribution of fitness effect of mutations), which both decrease the ER probability. On the contrary, increasing *r_max_* increases the ER probability through two effects. First, the size of the phenotypic space of resistance increases with *r_max_* (as in the SSWM regime, see Anciaux *et al*. 2018). Second a large *r_max_* counterbalances a high mutational load *nμ*/2 as can be seen in Eq.[2] (this latter effect is only captured in the WSSM approximation).

Figure 1 illustrates how the ER probability drops sharply with the decay rate *r_D_*, both in the SSWM regime (brown curves, *U* « *U*_C_, see legend) and in the WSSM regime (blue curves, *U* » *U_C_*). This qualitatively similar behavior is *a priori* due to common geometric constraints imposed by the FGM. The model of (Anciaux *et al*. 2018) and the present model, derived under complementary approximations (SSWM *versus* WSSM), capture the results of simulations in a different and complementary portion of the range of possible mutation rates: compare the blue (*U* = 10^2^ *U*_*c*_) vs. brown (*U* = 10^−2^ *U*_*c*_) curves to the dots in Figure 1. Higher mutation rates allows higher stress levels (larger *r_D_*) to be endured, but it is not their only effect, as we now detail.

### Non-monotonic relationship between ER probability and mutational parameters

In the following section, we investigate the effect of mutational parameters. Both the mutation rate *U* and the variance of mutational effects *λ* affect the system in a similar fashion through the composite parameter 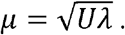 At small *t* (in Eq.[4]), an increase in *μ* speeds-up the early adaptive process, thus favoring rescue but also increases the ultimate mutation load, favoring extinction by lethal mutagenesis. These antagonistic effects of *μ* create a non-monotonic relationship between the rescue probability and mutational parameters. This is illustrated in Figure 2, which also shows how Eq.[5] (thick lines) captures this effect: not neglecting the parameter 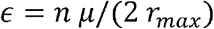 makes the relationship between *μ* and *P_R_* non-monotonic. For low stress *r_D_, P_R_* is maximal and approximately equal to 1 over a range of mutation rates (plateau Figure 2, see also Supplementary Figure 6 for higher stress values). Beyond this range, the rescue probability in Eq.[5] drops to 0 at 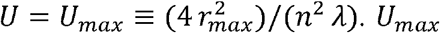 is the mutation rate beyond which certain extinction is enforced by lethal mutagenesis because the mutation load 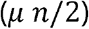 is larger than the maximal growth rate that can be reached in the stress (*r_max_*). Hence, even if ER allowed the population to invade the new environment, it could not generate a stable population once at mutation-selection balance.

Note that Eq.[6], which is only valid for intermediate *μ*, does not capture the decrease in *P_R_* close to *U* = *U_max_*. It does however capture the increase in *P_R_* as mutation rate increases, far below *U_max_*.

### Evolutionary dynamics from an initially polymorphic population (at mutation-selection balance)

The results presented in the previous figures illustrate rescue from *de novo* mutations. In the presence of additional standing genetic variance, rescue mutants can arise from *de novo* mutants, from pre-existing genotypes or from a combination of both. Figure 3 shows the qualitative similarity between the case with and without standing genetic variance, in their dependence on and, as observed in simulations and captured by Eqs.[5] and [6]. Indeed, Figure 3 confirms that the addition of standing genetic variance does not qualitatively modify the relationship between the rescue probability and stress intensity (, Figure 3a) or mutational parameters (here, Figure 3b). Note that the accuracy of Eq.[5] is lower for higher, where the continuous time approximations become less accurate to capture discrete time simulations (see Supplementary Figure 4). In the next sections, for the sake of clarity and simplicity, we will mainly discuss the scenario of ER from *de novo* mutations only, as the qualitative behaviors are similar with an extra contribution from standing variance.

**Figure 3:**
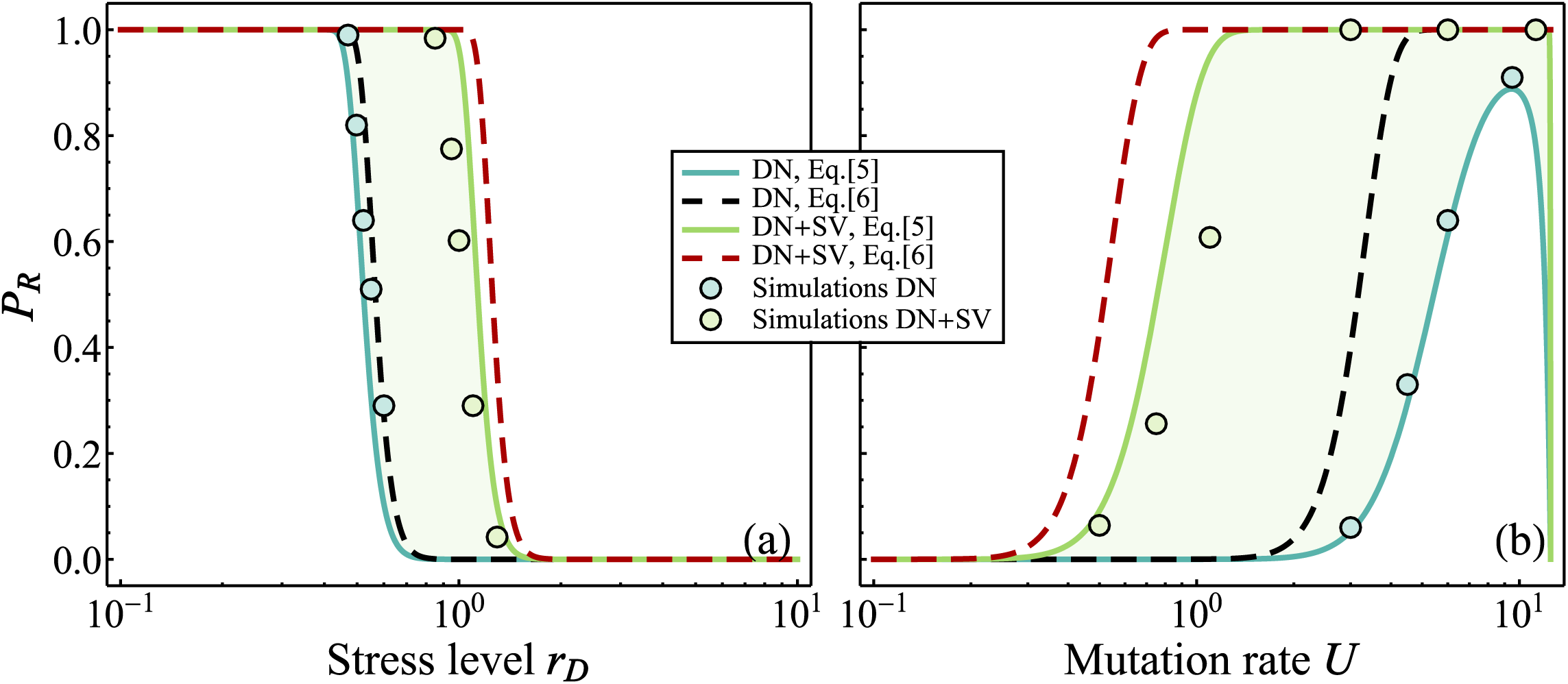
ER probabilities from *de novo* mutants only (blue: Eq.[5], black dashed line: Eq.[6]) or from both *de novo* and pre-existent mutants (green: Eq.[5], red dashed line: Eq.[6]) against the stress level (a) or the mutation rate (b). Dots show the results of simulations (started from 10 simulated populations at mutation-selection balance for the scenario). The shaded area corresponds to the contribution from the standing genetic variance to the rescue compared to *de novo* mutation. Other parameters are,, and. in panels (a) and in panel (b).

### Mutation window for ER

In the previous subsections, we have shown that the ER probability drops sharply with increasing stress and is maximal over a finite range of mutation rates, which we denote “mutation window” for ER. The “width” (range of mutation rates) and “height” (maximum of ER probability over the range) of this window strongly depends on stress.

#### Width of the window

To characterize the mutation window, its upper and lower bounds must be defined. The lower bound of the window (denoted *U*_*_) corresponds to the mutation rate at which the rescue probability rises to 1/2. Thus, this lower bound is only defined if the height of the window lies above or at 1/2 (i.e. max (*P_R_*) ≥ 1/2). The upper bound is set to the mutation rate *U_max_* beyond which certain extinction is enforced by lethal mutagenesis. The ER probability drops off very sharply close to *U_max_*, so that, approximatively, ER is only likely within the mutation window 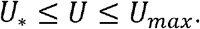 These two bounds are derived in Appendix I Eq. (A17):

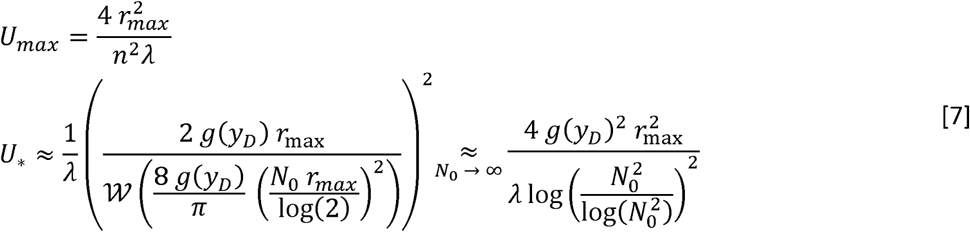

where *W*(.) is the Lambert *W* function, which converges to 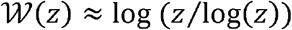 as (gets large (yielding the right hand side approximation above). Here, *g*(*y_D_*) is the function of stress intensity given in Eq.[6], which describes how stress intensity (*y_D_* = *r_D_/r_max_*) affects ER rates. Depending on the scenario, one uses 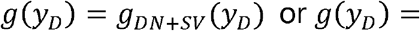 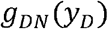 in the presence or absence of standing genetic variance, respectively (note that the window is always wider in the former case).

Figure 4 illustrates, for an initially clonal population, the sharpness of the transition from ER being almost certain to being highly unlikely (*P_R_* = 1 to *P_R_* = 0) as a function of *U* and *r_D_*. It also shows the accuracy of the approximation for *U*_*_ in Eq.[7] (dashed black line corresponding to the right hand side approximation of Eq.[7]), compared to its numerical estimation from Eq.[5] (color gradient). Supplementary Figure 5 shows a similar result for a population starting with standing genetic variance.

**Figure 4:**
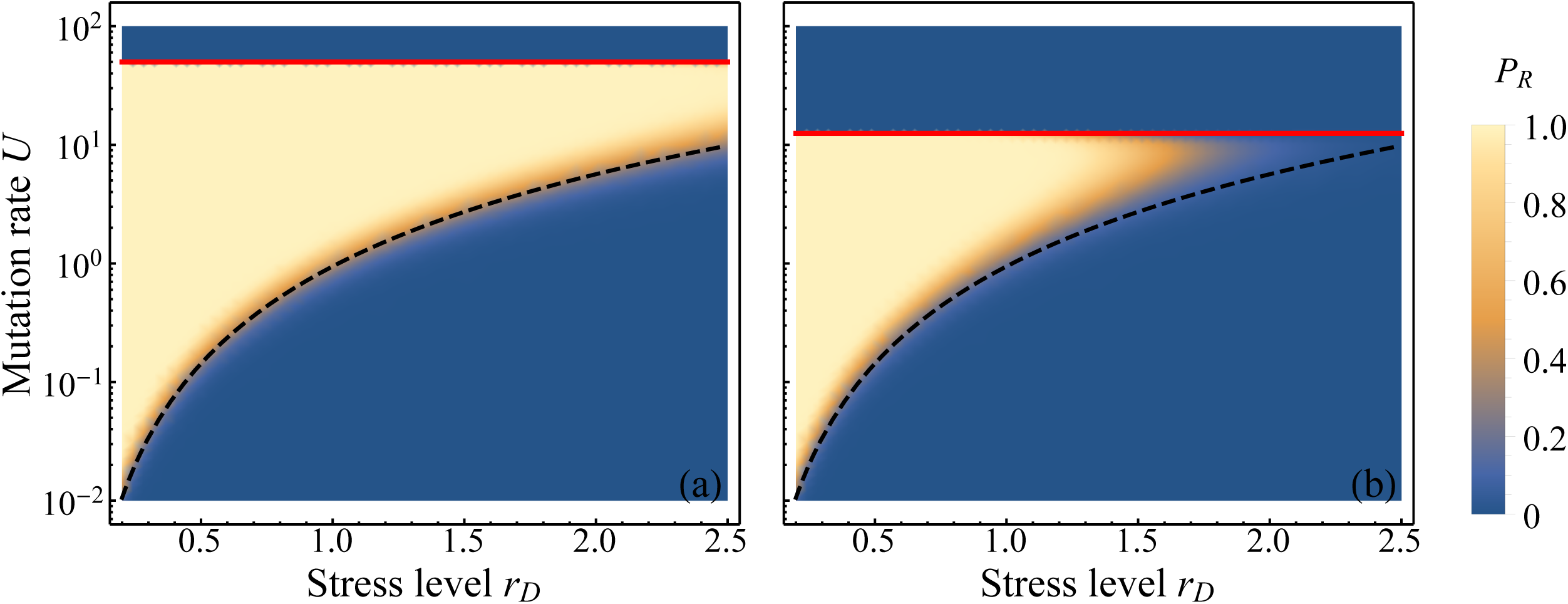
ER probabilities from *de novo* mutants only (Eq.[5]) for different values of *r_D_* and *U*. The color gradient gives the value of *P_R_* (see legend). The red straight line corresponds to *U* = *U_max_*(Eq.[7]) and the black dashed line corresponds to 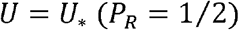 (right hand side approximation of Eq.[7]). For a given *U*, the ER probability drops sharply from *P_R_* = 1 (light yellow) to *P_R_* = 0 (blue), over a short increase in *r_D_*. For a given *r_D_, P_R_* rises sharply as increases about *U*_*_ and then drops sharply as increases around *U_max_*. Other parameters are *r_max_* = 1, *N*_0_ = 10^4^ and *λ* = 5 x 10^−3^. *n* = 4 in panel (a) and *n* = 8 in panel (b).

The upper bound *U_max_* is independent of initial conditions or stochasticity, as lethal mutagenesis depends on the deterministic equilibrium state of the population, once adapted to the stress. Therefore, *U_max_* does not depend on the presence or absence of initial standing variance, the decay rate imposed by the environmental change (*y_D_*) or the initial population size *N*_0_. On the contrary, the lower bound *U*_*_ depends on these factors as it is determined by the capacity of the population to transiently adapt to the new conditions. It shows, however, little dependence on dimensionality (*n*), as is also apparent in Figure 4, by the accuracy of Eq.[7], which is independent of *n*. Overall, the width of the mutation window where ER is likely decreases with increasing stress *r_D_* (Figure 4) and increases with initial population size *N*_0_(eq. [7]: *U*_*_ decreases with *N*_0_, and *U_max_* is unchanged). The width of the window decreases with dimensionality *n* and increases with the maximum fitness in the new environment *r_max_*, because both parameters affect the upper bound of the window, *U_max_* (eq. [7]). Lower dimensionality and higher maximum population growth rate reduce the mutation load, and thus allow persisting under higher mutation rates.

Finally, note that we focused on the effect of *U* here, but similar results could be obtained if *λ* was varied (as both parameters affect *P_R_* as a product 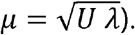 This is apparent in Eq.[7] where *U* and *λ* could be exchanged.

#### Height of the window and “Mutation proof extinction”

within the mutation window 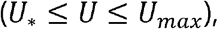 the ER probability *P_R_* rises above 50% and then drops back to zero. Yet, for more extreme stresses, it cannot even reach above 50% for any value of the mutation rate: the height of the mutation window lies below 1/2). When this height is low, extinction is ‘mutation proof’, in that it is highly likely whatever the mutation rate(s) *U* or the variance of mutational effects *λ* in the population. To illustrate this, we compute the maximum of the ER probability max(*P_R_*) when *U* varies from *U_C_* to *U_max_*, by numerically evaluating Eq.[5] over this range. Supplementary Figure 6 shows detailed profiles of ER probabilities against mutation rates (illustrating how max (*P_R_*) is found), in the presence or absence of initial standing variance. Figure 5 shows the maximum *P_R_* attainable as a function of *r_D_* and *N*_0_: it drops (transition from yellow to blue areas) with increasing stress *r_D_* and decreasing population size *N*_0_. In this example, a large part of the combinations of the two parameters *N*_0_ and *r_D_* correspond to max (*P_R_*) lower than 10% (blue area below the lower white dashed line in Figure 5). Therefore, for a given inoculum size *N*_0_, there is always a threshold of stress level beyond which ER is nearly impossible, whatever the mutation rates in the population.

**Figure 5:**
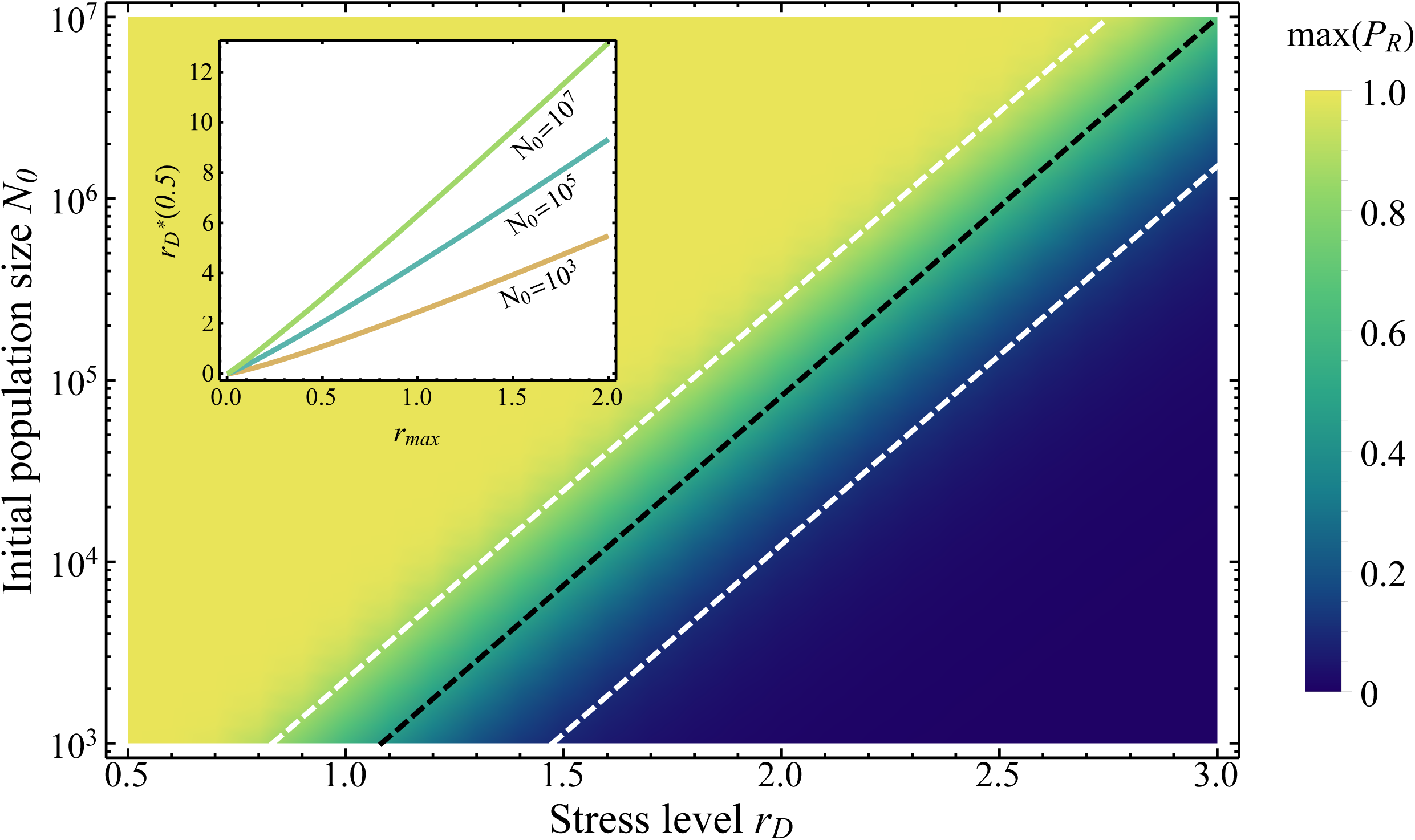
Maximum ER probability reached as is varied, for different values of and for a population with no initial polymorphism. The color gradient gives the value of this time (see legend). The black dashed line gives the value of (Eq.[8] with) and the white dashed lines the value of and. The maximum of the ER probability attainable (for all possible) drops sharply over a short range of increasing for a given, or over a short range of decreasing for a given. Other parameters are,, and. The inset panel shows how from Eq.[8] varies with and.

A closed form approximation can be obtained to describe this transition (detailed in Appendix I section VII): denote *r^*^_D_* the value of *r_D_* at which max(*P_R_*) = *p* for some *p* ∈ [0, 1]. We obtain the following simple expression for the threshold of level of stress beyond which max (*P_R_*) cannot exceed some level *p*, independently of *μ*;

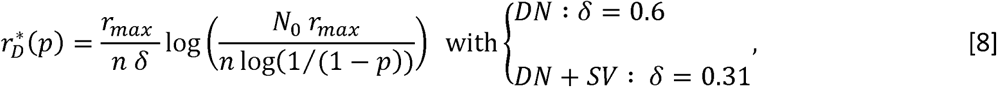

where the values of ϔ come from a curve-fitting procedure (detailed in Appendix I section VII). Setting *p* « 1 in Eq.[8] thus provides the stress level beyond which ER is very unlikely, whatever the mutation rate *U* or the variance of mutational effects *λ*. This means, in particular, that the evolution of higher mutation rates (via hypermutator strains) or higher variance of mutational effects (larger *λ*) would not allow the population to avoid extinction, when confronted to this stress level. The validity of the heuristic in Eq. [8] is illustrated in Figure 5, where we see that the dashed lines 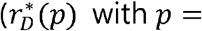 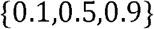 with p = { 0.1,0.5,0.99} see legend) accurately predict the transition from high to low values of max (*P_R_*), computed numerically from Eq.[5].

This whole argument applies to both *DN* and *DN* + *SV* scenarios (by choosing ϔ accordingly in Eq. [8]). Interestingly, we can also see that populations initially at mutation-selection balance (standing genetic variance) can withstand stresses twice larger than populations only adapting from *de novo* mutations (initially clonal).

#### Distribution of extinction times

From Eq.[4] we can derive the probability density of this distribution, in either of the two scenarios considered (purely clonal population or population at mutation-selection balance). We get:

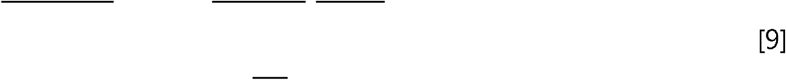

where the functions and depend on the scenario considered and are given explicitly in Eq.[5]. Figure 6 illustrates the accuracy of this result and how the distribution of extinction times varies with stress intensity and mutation rate. In spite of neglecting evolutionary stochasticity, Eq.[9] still captures the shape and scale of extinction time distributions, in the WSSM regime.

**Figure 6:**
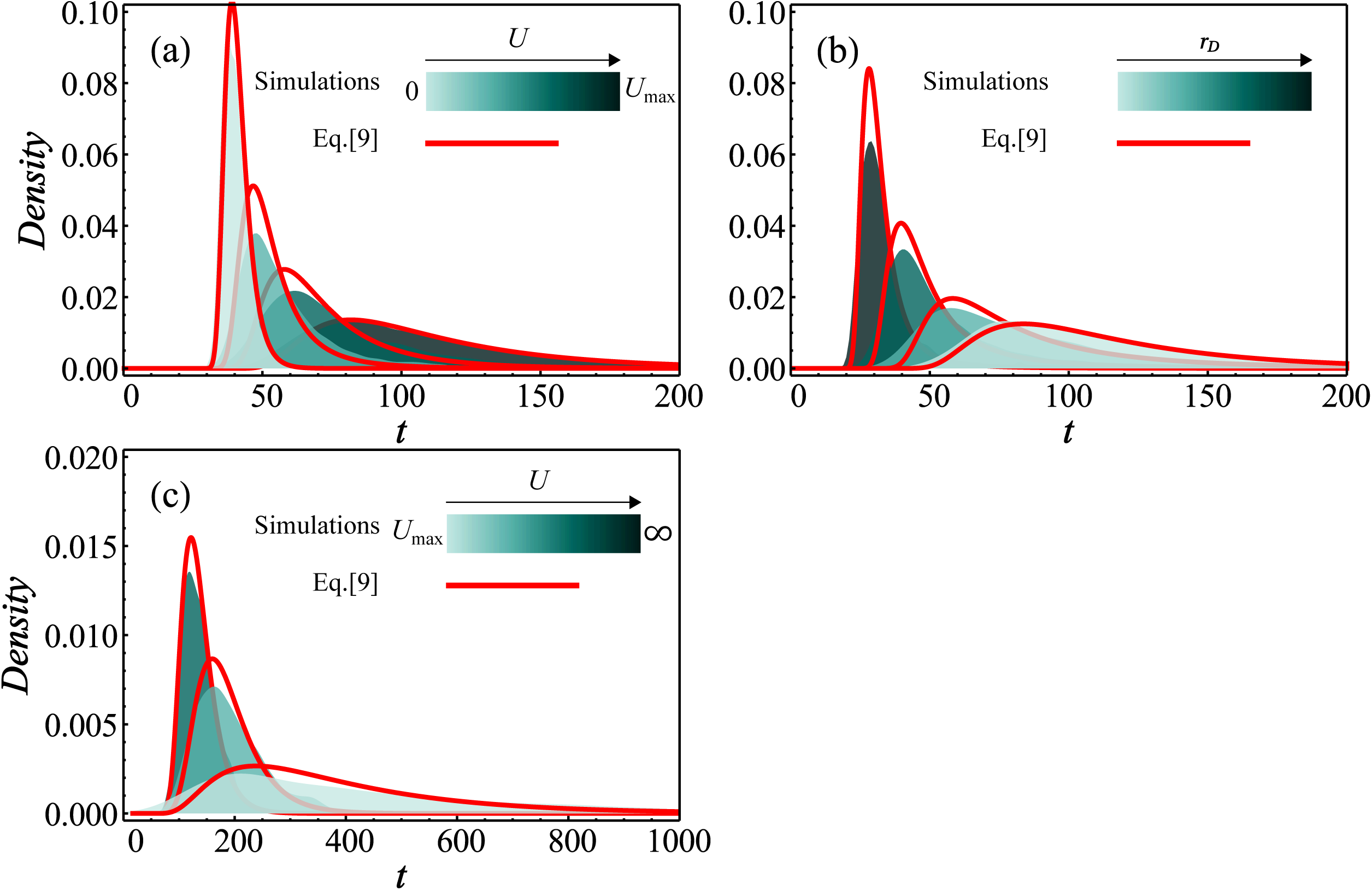
Density of the extinction probability dynamics for different values of (panel (a) and (c)) and (panel (b)). The distributions of extinction times from simulations started with an isogenic population are shown by shaded histograms, with the corresponding theory (Eq.[9]) given by the plain red lines. The color gradient corresponds to increasing levels of *r_D_* (panel (b)) or of *U* (lower range in panel (a) and higher range in panel (c), as indicated on the legend). Each simulated distributions is drawn from 1000 extinct populations among a sufficient number of replicates to observe these 1000 extinctions. The mutation rates covered in panel (a) are *U* = {0; 4 *U_C_*; 8 *U_C_*; 14 *U_C_*} and in panel (c) are *U* = {2 *U_max_*; 1.5 *U_max_*; 1.1 *U_max_*} and in both panel the decay rate is *r_D_* = 0.28. The decay rate covered in panel (b) are {0.285; 0.33; 0.4; 0.59} and the mutation rate is *U* = 15 *U_c_*. Other parameters are *N*_0_ = 10^5^, *n* = 4, *r_max_* = 0.1 and *λ* = 5 x 10^−3^.

Figure 6b shows that decreasing the stress level (*r_D_*) increases the mean persistence time of the population and also increases the variance of this duration. This behavior could be seen as a mere scaling: even in the absence of evolution the population takes longer time to become extinct with a smaller *r_D_*. On the contrary, increasing the mutation rate, keeping it below the lethal mutagenesis threshold (0 ≤ *U* ≤ *U_max_*, Figure 6a), increases the mean and the variance of the persistence time. This likely stems from subcritical mutations (0 > *r* > *r_D_*, beneficial but not resistant) that can transiently invade the population, thus delaying its extinction. However, beyond the lethal mutagenesis threshold (*U* > *U_max_*, Figure 6c), the trend is reversed: extinctions (which are always certain then) occur faster at high mutation rates (panel c). Therefore, even in those cases where ER probabilities are uninformative (*P_R_* = 0), the distribution of extinction times conveys important information on the underlying adaptive or maladaptive dynamics.

## Discussion

We investigated the effect of an abrupt environmental change on the persistence of asexual populations with a large mutational input of genetic variance (WSSM regime), adapting either from *de novo* mutations arising after the environmental change (*DN* scenario) or from both *de novo* and pre-existing mutations (*DN* + *SV* scenario). In a previous study (Anciaux *et al*. 2018), we studied evolutionary rescue when considering adaptation over a phenotype-fitness landscape (FGM), which implies pervasive epistasis between multiple mutations and imposes a relationship between the initial decay rate of the population, the proportion and growth rate of resistance alleles, and their selective cost in the ancestral (before the stress) environment. However, we assumed that rescue resulted from rare mutations with strong effects (SSWM). The key contributions of the present model, building on our previous work, are to (i) allow for the cumulative effect of multiple mutations and (ii) provide insights into the distribution of extinction times in the presence of an evolutionary response.

### Single step (SSWM regime) vs. multiple step (WSSM) regimes in ER

In spite of its complexity, the ER process in high mutation rate regimes can readily be captured by simple analytic approximations (Eqs.[5] and [6]), which neglect evolutionary stochasticity and only account for demographic stochasticity. Overall, the SSWM and WSSM approximations roughly capture the ER process in complementary domains of the mutation rate spectrum (Figures 1-3).

This approach shows how multiple mutations allow withstanding higher stress than what the single step approximation (SSWM in Anciaux *et al*. 2018) predicts (Figures 1 and 3). However, this is only true for intermediate mutation rates: a further increase in mutation rate ultimately shifts the system to a lethal mutagenesis regime (Figures 2 to 4). Indeed, the dependence between the ER probability and the mutation rate is not monotonic. The model shows an optimal mutation rate for the ER probability, at which the maximal ER probability may be less than 1 (depending on the stress, Figure 5). Beyond this rate, the ER probability drops down, to some point (*U_max_*, Eq. [7]) where the mutation load is so large that absolute fitness is negative at mutation selection balance. This non-monotonic dependence reflects the continuum between ER and lethal mutagenesis along a gradient of mutation rates.

### Similarities to existing ER models

Some of our key previous findings (Anciaux *et al*. 2018), regarding how ER depends on the parameters of the fitness landscape (FGM here) are still valid in the more polymorphic WSSM regime. The main common features are (i) the sharp decrease of ER probabilities with stress (decay rate *r_D_*), (ii) their log-linear increase with initial population size *N*_0_, (iii) the fact that standing variance allows withstanding higher stress, (vi) the limited effect of dimensionality *n* (for mutation rates far from lethal mutagenesis, eq. [6]). The effect of initial population size (*P_R_* = 1 − exp (− N_0_ *ω*) eq.[5]) is exactly the same as in previous models where ER stemmed from single mutants (SSWM regime Orr and Unckless 2008, 2014; Martin *et al*. 2013; Anciaux *et al*. 2018). This is expected of any model ignoring interactions between individuals, be it evolutionary (e.g. sexual reproduction or frequency-dependent selection) or demographic (e.g. density-dependence): each of the *N*_0_ lineages initially present contributes independently to ER (with some rate *ω* per individual). Decay rate has a broadly similar (but quantitatively different) effect in previous ER models not based on a fitness landscape. The other parameters (*r_max_*, *n*, *λ*) are not defined outside the FGM. More generally, the key implications of considering the FGM to model ER are detailed more thoroughly in Anciaux *et al*. (2018).

### Experimental test and parametrization

To experimentally test the predictions, the assumptions of the WSSM regime *a priori* imply the need to use organisms with relatively high mutation rates, such as viruses, highly mutating strains of bacteria or possibly cancer cells. How large should the mutational parameters be for the WSSM to apply? Empirically mutation effects and rates, at least in microbes, are typically scaled by growth rates. In our model, if this growth rate is the maximal growth rate (that of the wild type adapted to the lab environment), and assuming the lab environment and the stressful environment have the same *r_max_*, then, these scaled parameters would correspond to s 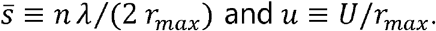. If *r_max_* is approximately equal to a birth rate (i.e. the optimal ‘wild-type’ has a small death rate in the lab environment) 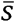 is akin to a mean mutation effect per division, while *u* is akin to a mutation rate per division. Expressed in these parameters, the WSSM approximation (Eq.[5]) applies whenever 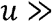 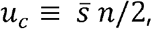, lethal mutagenesis occurs when 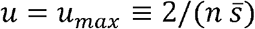 and the intermediate mutation weak selection approximation (Eq.[6]) is valid when 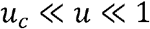 (see the section weak selection intermediate mutation approximation from the results). Estimates of dimensionality *n* are typically not very large based on mutation fitness effects analyzed under the FGM (Martin and Lenormand 2006; Perfeito *et al*. 2014). Therefore, based on this very rough analysis, the WSSM ER model proposed here would apply in strains with mutation rates *u* higher than mutation effects 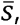 measured in the same per division time units.

For a proper experimental test of the predictions, the parameters 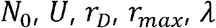 and *n* must be measured, in the stressful environment(s) where ER is studied. The methods and challenges in estimating these parameters are discussed in Anciaux *et al*. (2018). Note however that (i) the WSSM results only depend on the product 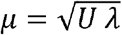 and that (ii) an error on the dimensionality may not be critical given its relatively small effect (except in setting *U_C_* above which the model applies). The composite parameter *μ* can be estimated directly from mutation accumulation (MA) experiments as 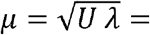 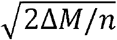 where 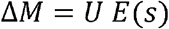 is the change in mean fitness per unit time in the MA experiment 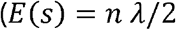 is the mean selection coefficient of random mutations).

Finally, if the model is valid, fitting an observed distribution of extinction times (Figure 6) might provide estimates of *r_D_, r_max_*, *n* and possibly *μ* (note however that large or low values of *μ* > *μ_C_* could produce similar distributions, see Figure 6). This would allow using not only the information from rescued populations but that from extinct ones. Such empirical test would require fine-scale time series of the population size or at least the extinction status over time.

### Treatment against pathogens, hyper-mutators and lethal mutagenesis

Our model as well as previous ones all suggest that the effectiveness of a given treatment depends on the mutation rate of the organism. Polymorphism for mutation rate and invasion of hyper-mutator genotypes are thus potentially important issues for treatments against pathogens. However, our results suggest that a sufficiently strong stress could be effective in spite of hyper-mutator evolution. Indeed, because of the lethal mutagenesis effect (Figure 2), ER is only possible within a mutation rate window: a hyper-mutator would have to hit this window to be advantageous, and the width of the window narrows with increased stress (Figure 4). At sufficiently higher stress levels, ER is unlikely whatever the mutation rate (Figure 5), making these strong treatments robust to hyper-mutator evolution. Whether this pattern is confirmed empirically and whether the required treatment levels are then not too harmful for the treated subject remain open questions. Note also that the same line of argument could be used, not for mutation rate evolution, but for the evolution of the variance of mutational effects, as the end result depends on the product 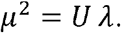

Our model covers the continuum from stress induced extinction to extinction induced by lethal mutagenesis. The latter might be an option, especially for organisms for which no “stress treatment” exists, or whose high mutation rates (above *U*_*_ in our model) allows them to withstand even strong stresses. Our results indeed confirm that increasing the mutation rate in this context (above *U_max_*) will allow to completely eliminate the population. The addition of a stressor might also help in the process, as has been suggested before (Pariente *et al*. 2001, 2003, 2005): indeed, the ER probability does drop faster with increasing *U* (as we approach *U_max_*) in the presence of a strong stress (with high *y_D_*), see Supplementary Figure 6.

### Treatment duration issues

One may also wonder how long a treatment must last (be it by stress effect or by lethal mutagenesis) for it to be efficient. From the distribution of extinction times (Figure 6), it is possible to predict the duration of the treatment needed to get rid of, say, 99% of the pathogen populations. Strong stresses (dark blue histogram in Figure 6b) tend to both decrease the overall ER probability and shrink the distribution of times to extinction. This means they are more efficient overall and require shorter treatment durations, with less risk associated with imperfect treatment compliance. However, when the mutation rate increases towards *U_max_* (treatment by lethal mutagenesis), although extinction becomes highly likely (*P_R_* → 0, Figure 2), extinction times get more variable and the mode of the distribution is higher (Figure 6a). If the mutation rate increases beyond *U_max_*, the opposite pattern is observed: the distribution of extinction times shrinks as *U* increases further away from *U_max_*(Figure 6c). Hence, a treatment by lethal mutagenesis, even if guaranteeing total extinction after infinite time, may need to be applied for a long time to significantly decrease ER probability (if the resulting *U* is close to *U_max_*). Thus, observed distributions of extinction times hold valuable information.

### Limits of the model

Obviously, the different results and applications described above are limited by the hypotheses of the model.

First, the model ignores density (discussed in Anciaux *et al*. 2018) or frequency-dependent effects. Hence, our model cannot describe “competitive release” effects which have been discussed in the context of antibiotic resistance (Read *et al*. 2011; Day and Read 2016), whereby higher stresses, by eliminating sensitive genotypes more rapidly, can “release” limiting resources for resistant ones, and ultimately increase ER probability. However, we do not expect *a priori* the present model to show such a decrease in ER probability at higher stress levels. In the FGM in particular, given the very sharp decay in *P_R_* with stress level *r_D_*, it is highly likely that density-dependence (hence room for competitive release) will only mitigate the result and that ER probability will still decrease at higher stress. However, this must be studied quantitatively in a dedicated model, which would account for density-dependence.

The approximation of the demographic dynamics by a Feller diffusion also imposes that demographic variations remain smooth, which may be inaccurate, e.g. for some viruses showing occasional large burst events. The Feller diffusion can also fail to predict discrete time demography (used here in the simulations) at high growth rates per generation 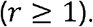

Second, in the demographic dynamics, we also make the approximation that σ is roughly constant across genotypes and thus constant over time. This approximation can be accurate if the mutant growth rates are typically small compared to their reproductive variance *r_i_* « *σ_i_*, which is valid as long as 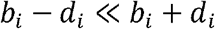 (with *b_i_* (resp. *d_i_*) the birth rate (resp. the death rate) of each genotype; discussed in Martin *et al*. 2013). However if mutations affect substantially but dissimilarly the birth and death rates this approximation might fail. This could be handled by considering the deterministic time dynamics of the mean reproductive variance 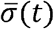 in addition to that of the mean growth rate 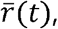 in the inhomogeneous Feller diffusion.s

Third our results assume a sharp change in the environment, which does not reflect all forms of stresses, including in the context of sudden treatment (e.g. antibiotics can have complex pharmacokinetic patterns over time Regoes *et al*. 2004).

## Conclusion

Recently, lethal mutagenesis treatments have received renewed interest for their ability to provide alternative treatments against viruses resistant to other type of drugs (Jiang *et al*. 2016; Escribano-Romero *et al*. 2017). The present model attempts to provide a framework to predict the efficacy of such treatments. The strategy used here of coupling deterministic evolutionary trajectories with stochastic demographic dynamics could in principle be applied to other models, to cope with a large input of mutations affecting the ER process. We hope that future experimental tests will evaluate its accuracy and potential to tackle various pressing applied issues.

## Supporting information

Matlab code for curve fitting

Mathematica Notebook with Analytical derivations, Figures code and Simulation code

