## Appendix I for "Population persistence under high mutation rate: from evolutionary rescue to lethal mutagenesis"

**Appendix I : Mathematical derivations and approximations**

In the following, we use a supplementary notation to simplify the mathematical derivations: $\theta=n/2$.

**I. General WSSM approximation to ER.**

In the present model, we approximate the stochastic dynamics of each lineage by a Feller diffusion (Feller 1951) or continuous branching CB-process (Lambert 2008), with parameters$\{r_{i},\sigma_{i}\}$ for lineage$i$. We ignore any density or frequency dependence and assume that all lineages have similar stochastic reproductive variance ($\sigma_{i}\approx\sigma$). The resulting total population size$N_{t}$ (cumulating all lineage that co-segregate) can then also be approximated by a CB process, which follows the stochastic differential equation:

$$dN_{t}=\bar{r}_{t} N_{t} dt+\sqrt{{\sigma N}_{t}} dB_{t} ,$$

(A1)

where$B_{t}$ is a Weiner process,$\bar{r}_{t}=1/N_{t}\sum_{i=1}^{N_{t}} r_{i}$ is the mean growth rate of all lineages present at time$t$ and$\sigma$ is the common stochastic reproductive variance of all lineages. In the WSSM regime (i.e. when$U\gg U_{c}=\theta^{2} \lambda$), we can ignore the evolutionary stochasticity introduced by mutation and drift as a first approximation. Then, the mean growth rate $\bar{r}_{t}\approx\left\langle\bar{r}_{t} \right\rangle$ (expectation over replicates denoted by$\left\langle. \right\rangle$) is approximately deterministic and given by the WSSM results in (Martin and Roques 2016) for the FGM. The probability of such a time inhomogeneous CB process to be extinct by time$t$ is given by (Bansaye and Simatos 2015):

$$P_{\mathrm{ext}}\left( t \right)=\exp\left( -\frac{2N_{0}}{\psi_{t}} \right) ,$$

(A2)

where$\psi_{t}=\sigma\int_{0}^{t} exp(-\rho(v))dv$ and$\rho(u)=\int_{0}^{u} \left\langle\bar{r}_{v} \right\rangle dv$. The probability of ER is the complementary probability of not being extinct over infinite time, namely$P_{R}=1-P_{\mathrm{ext}}(\infty)$.For consistency with the main text, $\sigma\approx1$ in the following.

**II. Explicit expression for a population initially clonal**

Define $r_{D}$ the decay rate of the initial clone, and$r_{max}$ the maximal attainable growth rate (that of a genotype optimal in the stressful environment). In this case, the mean fitness trajectory, relative to the optimal genotype ($\left\langle\bar{m}_{t} \right\rangle=\left\langle\bar{r}_{t} \right\rangle-r_{max}$), under the WSSM approximation, for an initially clonal population is (eq. (12) in (Martin and Roques 2016)

$$\left\langle\bar{m}_{t} \right\rangle=m_{0} {(sech\left( \mu t \right))}^{2}-\theta\mu\tanh\left( \mu t \right) ,$$

(A3)

where $\mu=\sqrt{U \lambda}$ and$\theta=n/2$, $\mathrm{sech}(.)$ and $\tanh(.)$ are the hyperbolic secant and tangent functions and $m_{0}$ is the fitness difference between the original clone and the optimal genotype. Now, we require the absolute mean fitness trajectory (mean growth rate), which is obtained by noting that, by definition,$\left\langle\bar{m}_{t} \right\rangle=\left\langle\bar{r}_{t} \right\rangle-r_{max}$ and $m_{0}=-r_{D}-r_{max}$. Denoting$y_{D}=r_{D}/r_{max}$,$\epsilon=\theta\mu/r_{max}$ we obtain:

$$\left\langle\bar{r}_{t} \right\rangle=\frac{\theta\mu}{\epsilon} (1-\left( y_{D}+1 \right)\mathrm{sech} \left( \mu t \right)^{2}-\epsilon tanh(\mu t)) .$$

(A4)

The integral $\rho_{t}=\int_{0}^{t} \left\langle\bar{r}_{v} \right\rangle dv$of this growth rate over time can be expressed in compact form by using the change of variable$\tau=\mu t$, yielding

$$\begin{matrix} \rho_{t}=\frac{\theta}{\epsilon}f(\tau)-\theta\log\left( h\left( \tau\right) \right) \\ h\left( \tau\right)=cosh(\tau) \\ f\left( \tau\right)=\tau-\left( 1+y_{D} \right)\tanh\left( \tau\right) \end{matrix}.$$

(A5)

Finally, the same change of variable ($dt=\mu d\tau$) can also be used to express the indefinite integral that determines extinction probabilities, yielding the rate of ER per lineage present at the onset of stress:

$$\begin{matrix} \omega^{DN}=-\frac{\log P_{\mathrm{ext}}\left( \infty\right)}{N_{0}}=\frac{2}{\int_{0}^{\infty} exp(-\rho_{u}) du} \\ \omega^{DN}={2 \mu}/{(\int_{0}^{\infty} h\left( \tau\right)^{\theta} exp(-\frac{\theta}{\epsilon} f\left( \tau\right))ⅆ\tau)} \end{matrix} .$$

(A6)

**III. Laplace approximations for small**$\boldsymbol{\epsilon}$**, for a population initially clonal**

Eq.(A6) is fully analytic but yields no explicit expression, which would be useful to get a more intuitive grasp of how each parameter affects the ER probability. When the mutation rate and effects are small enough that the load$\theta\mu$ is small relative to the maximal growth rate$r_{max}$ (so that we are well away from lethal mutagenesis), then$\epsilon\ll1$. This means that $\theta/\epsilon\gg1$ (with $\theta\geq1/2$) and the integral in Eq.(A6) is amenable to the Laplace Approximation (as$h\left( \tau\right)$ is monotonic over$\tau\in\mathbb{R}^{+}$). This approximation can be formulated as follows :

$$\int_{0}^{\infty} h\left( \tau\right)^{\theta} exp(-\frac{\theta}{\epsilon} f\left( \tau\right))ⅆ\tau\underset{\epsilon\to0}{\approx}\sqrt{\frac{2 \pi\epsilon}{\theta f\text{''}\left( \tau_{0} \right)}} h\left( \tau_{0} \right)^{\theta} exp(-\frac{\theta}{\epsilon} f\left( \tau_{0} \right))$$

(A7)

where $\tau_{0}=\cosh^{-1} (\sqrt{1+y_{D}})$ is the unique minimum of $f\left( . \right)$ over $\tau\in\mathbb{R}^{+}$. The Laplace approximation in Eq.(A7) then yields a fully explicit expression for the rate of ER per lineage present at the onset of stress defined in Eq.(A6). Rewriting in terms of the original parameters ($\theta/\epsilon= r_{max}/\mu$), we get

$$\begin{matrix} \omega^{DN}\underset{\epsilon\to0}{\approx}2 \sqrt{\frac{r_{max} \mu}{\pi}}\exp\left( -\frac{r_{max}}{\mu} \gamma\left( y_{D} \right) \right) \\ \gamma\left( y_{D} \right)=\sqrt{y_{D}\left( 1+y_{D} \right)}-\cosh^{-1}\left( \sqrt{1+y_{D}} \right)+\epsilon\left( \left( 1+\frac{1}{2 \theta} \right)\frac{\log\left( 1+y_{D} \right)}{2}-\frac{1}{\theta}\frac{\log\left( y_{D} \right)}{4} \right) \end{matrix} .$$

(A8)

The accuracy of this expression is illustrated in **Supplementary Figure 1**.

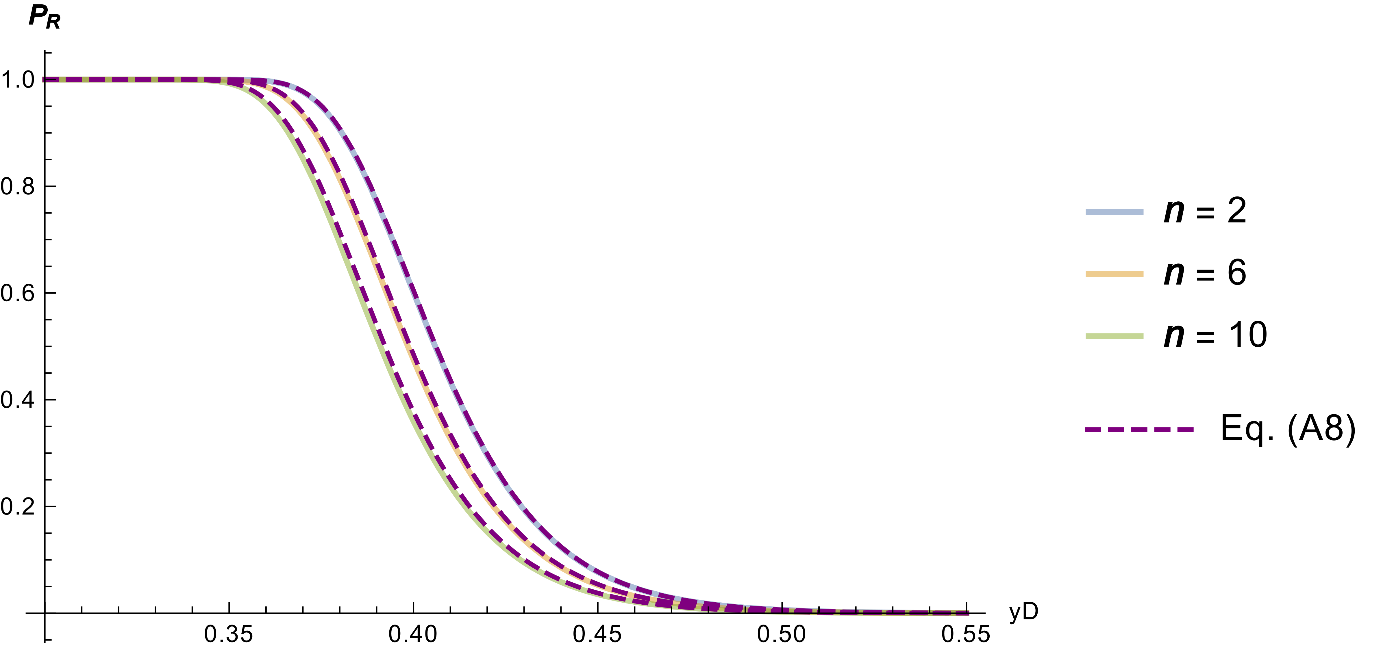

***Supplementary Figure 1:*** $P_{R}$ *as a function of*$y_{D}$ *for three values of the dimensionality (given in legend), as computed from the ‘exact’ Eq.(A6) (solid colored lines) vs. Laplace Approximation Eq.(A8) (dashed purple lines). Parameters are*$r_{max}=1,\lambda=5.{10}^{-4}, U=0.25,N_{0}={10}^{7}$*.*

In fact, unless we consider narrow range of stress variation, as in **Supplementary Figure 1**, it appears that, in the limit$\epsilon\to0$,$P_{R}$ shows limited dependency on dimensionality, provided that it remain limited ($n$ varies by five-fold above). This is confirmed by further simplifying Eq.(A8) to produce a rough but reasonably accurate approximation, in a range that is a priori of most biological relevance, say for $y_{D}>0.1$, i.e. not-too mild a stress. In this range both${\log\left( 1+y_{D} \right)}/2$ and ${\log\left( y_{D} \right)}/4$ are of similar or smaller order than$g\left( y_{D} \right)=\sqrt{y_{D}(1+y_{D})}-\cosh^{-1} \left( \sqrt{1+y_{D}} \right)$. Therefore, as$\epsilon\to0$ (as we assume here), the right hand factor in$\gamma(y_{D})$, proportional to$\epsilon$, becomes negligible, relative to the left hand term, and $\gamma\left( y_{D} \right)\approx g(y_{D})$. Eq. (A8) thus simplifies to the expression given in Eq.(6) for *de novo* rescue:

$$\begin{matrix} \begin{matrix} \omega^{DN}\underset{\begin{matrix} \epsilon\to0 \\ y_{D} \geq0.1 \end{matrix}}{\approx}2 \sqrt{\frac{r_{max} \mu}{\pi}}\exp\left( -\frac{r_{max}}{\mu} g\left( y_{D} \right) \right) \\ g\left( y_{D} \right)=\sqrt{y_{D}(1+y_{D})}-\cosh^{-1} \left( \sqrt{1+y_{D}} \right) \end{matrix} \end{matrix} .$$

(A9)

This simpler approximation is less precise but still provides a good order of magnitude, when$\epsilon$ is small enough. **Supplementary Figure 2** illustrates its accuracy and the fact that the ER rate is indeed roughly independent of dimensionality $\theta$ in this parameter range : Eq.(A9) captures the order of magnitude of stress at which $P{}_{R}$ drop from $1$ to $0$.

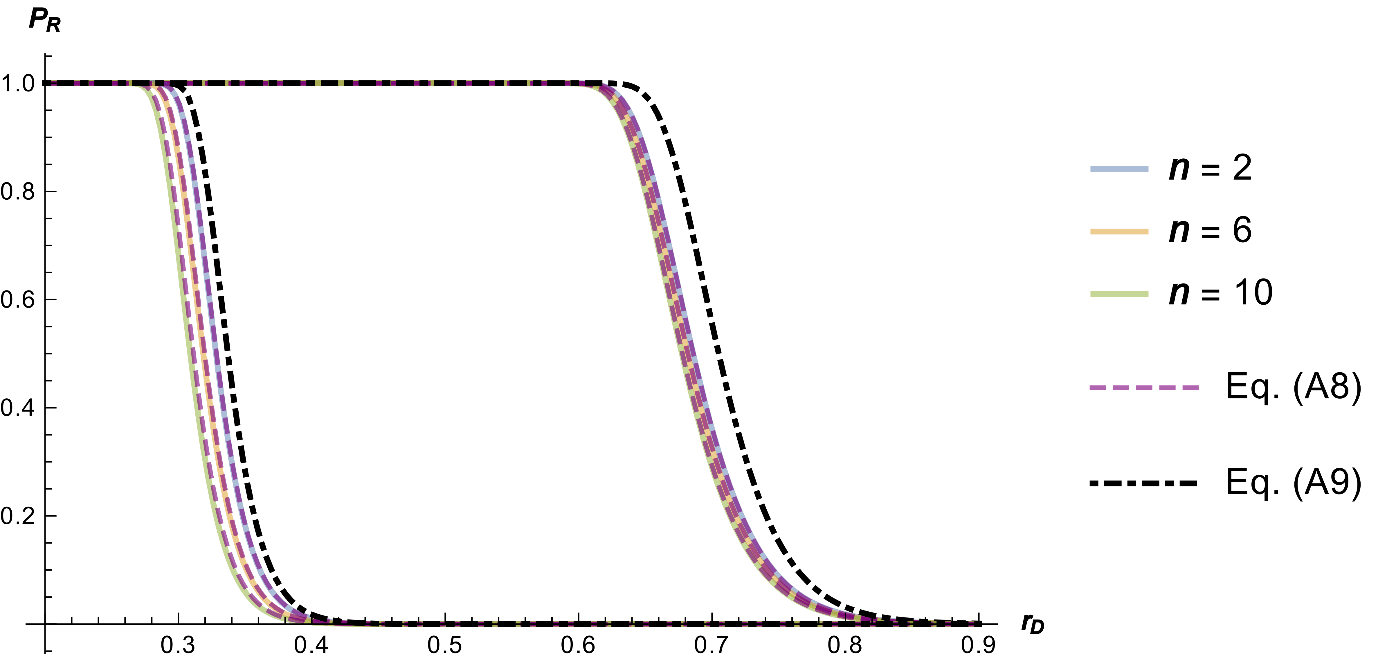

$$r_{max}=0.5$$

$$r_{max}=5$$

***Supplementary Figure 2:*** *Same as* ***Supplementary Figure 1*** *(same parameters, except*$r_{max}$*, indicated on the graph), this time as a function of*$r_{D}$ *and compared to both Eqs.(A8) (purple dashed lines) and (A9) (black dotdashed lines).*

**IV. Explicit expression for a population initially at mutation-selection balance**

This time we assume initially mutation-selection balance, then a shift in optimum occurs without any change in$U$ or$\lambda$: adaptation is driven by both standing variance and de novo mutations. Under the WSSM approximation (corresponding to our assumption here), the mutation selection balance in an asexual population under the isotropic FGM corresponds to a simple Gaussian distribution of phenotypes, centered on the optimum (in the environment where the equilibrium sets) and a variance $\mu$ per trait:$\mathbf{z} \sim N(\boldsymbol{0},\mu\mathbf{I}_{n})$. Once this optimum is shifted in the new environment, the phenotypic distribution progressively shifts (retaining the same normality and variance) towards the new optimum. This yields the following dynamics for the mean growth rate (based on the corresponding WSSM approximation in eq. (13) in (Martin and Roques 2016), with the same parameterization as for Eq.(A4)) we get

$$\left\langle\bar{r}_{t} \right\rangle=r_{max}(1-exp(-2 \mu t)(1+y_{D})-\epsilon) ,$$

(A10)

which then yields the corresponding expression for$\rho_{t}$ (same form as Eq.(A5)):

$$\begin{matrix} \rho_{t}=\frac{\theta}{\epsilon}f\left( \tau\right)-\theta log(h\left( \tau\right)) \\ f\left( \tau\right)=\tau-{\left( 1+y_{D} \right) \left( 1-exp(-2\tau) \right)}/2 \\ h\left( \tau\right)=exp(\tau) \end{matrix} .$$

(A11)

The rate of ER takes a similar form as in the DN scenario (Eq.(A6)), but this time an explicit exact expression is found:

$$\begin{matrix} \omega^{DN+SV}=- \log P_{\mathrm{ext}}\left( \infty\right)/N_{0}=\frac{2}{\int_{0}^{\infty} exp(-\rho_{u}) du}=2 \mu/\int_{0}^{\infty} h\left( \tau\right)^{\theta}exp(-\frac{\theta}{\epsilon} f\left( \tau\right))d\tau\\ \omega^{DN+SV}=4 \mu\frac{exp(- \xi) \xi^{\beta}}{\Gamma(\beta)-\Gamma(\beta,\xi)} \end{matrix}$$

(A12)

Where $\xi={\left( 1+y_{D} \right) \theta}/{(2 \epsilon)}$ and$\beta={\left( 1-\epsilon\right) \theta}/{(2 \epsilon)}$, and $\Gamma(.)$ and $\Gamma(.,.)$ are respectively the Euler’s gamma function and the incomplete gamma function.

**V. Laplace approximations for small**$\boldsymbol{\epsilon}$**, for a population initially at mutation-selection balance**

Although this is already explicit, a Laplace approximation can be used to get a simpler result, again away from the lethal mutagenesis regime ($\epsilon\ll1$). The function$f(.)$ has a unique minimum at$\tau_{0}=\log\left( \sqrt{1+y_{D}} \right)$ and the resulting Laplace approximation for the rate of ER is of the same form as Eq.(A8):

$$\begin{matrix} \begin{matrix} \omega^{DN+SV}\underset{\epsilon\to0}{\approx}2 \sqrt{\frac{r_{max} \mu}{\pi}}\exp\left( -\frac{r_{max}}{\mu} \gamma\left( y_{D} \right) \right) \\ \gamma\left( y_{D} \right)=\frac{y_{D}-(1-\epsilon)\log\left( 1+y_{D} \right)}{2} \end{matrix} \end{matrix} ,$$

(A13)

From which, as$\epsilon\ll1$, arises the same expression as Eq.(A9) with a different function$g(.)$:

$$\begin{matrix} \begin{matrix} \omega^{DN+SV}\underset{\epsilon\to0}{\approx}2 \sqrt{\frac{r_{max} \mu}{\pi}}\exp\left( -\frac{r_{max}}{\mu} g\left( y_{D} \right) \right) \\ g\left( y_{D} \right)=(y_{D}-\log\left( 1+y_{D} \right))/2 \end{matrix} \end{matrix} .$$

(A14)

The accuracy of this approximation is illustrated in **Supplementary Figure 3** below. Note also how the drop in ER probability occurs at higher stress levels here than with purely *de novo* mutation (in **Supplementary Figure 2**).

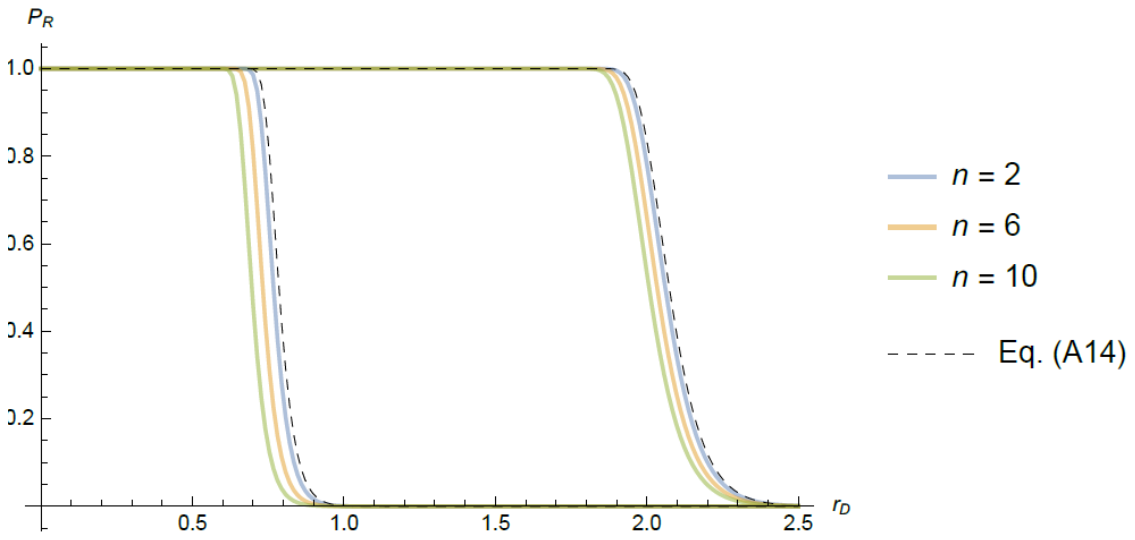

$$r_{max}=5$$

$$r_{max}=0.5$$

***Supplementary Figure 3:*** *Same as* ***Supplementary Figure 2*** *(same parameters) with standing variance plus de novo mutation computed from the ‘exact’ Eq.(A12) (solid colored lines), and the corresponding approximation Eq.(A14) (black dashed line).*

The accuracy shown in the main text between Eq.[5] (Eq.A(12)) and the simulations decrease when the growth rates of the individuals in the population increase (in absolute value). Indeed, when growth rates are too low or high $(\left| r \right|\geq1)$, the continuous time approximation used in our model fails to predict accurately the discrete time dynamic of the population size and therefore the probability of rescue given by our simulations. This is illustrated in **Supplementary Figure 4** where we can see that increasing $r_{max}$ leads to an underestimation of the rescue probability predicted by the simulations. Moreover, at higher $U$ the ER probability drop from highly likely to highly unlikely at larger $r_{D}$, of order $1$ or more, for which the discrepancies between continuous time and discrete time dynamics mentioned above increase, as shown in **Supplementary Figure 4.**

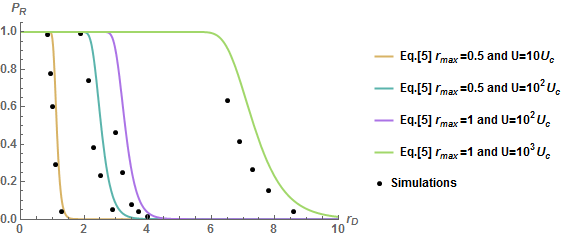

***Supplementary figure 4:*** *Accuracy of the predictions of ER probabilities for DN+SV scenario (blue: Eq.[5]) for two values of* $r_{max}$ *and two values of* $U$ *against simulations (see legend). Note that the green line and the corresponding simulations are the same as in Figure 3a. Other parameters are* $N_{0}={10}^{5}$*,* $n=4$*,* $\lambda={5.10}^{-3}$*.*

**VI. Width of the mutation window of ER**

We now wish to evaluate the mutation rate at which ER switches from very likely to very unlikely. We know that the rate of ER drops sharply at some upper threshold mutation rate, due to lethal mutagenesis effects (i.e. when the mutation load $\mu\theta=r_{max})$. This implies a maximal mutation rate $U_{max}=r_{max}^{2}/(\theta^{2}\lambda)$, above which lethal mutagenesis leads to certain extinction of the population.

Here, we are interested in the parameter range, far below lethal mutagenesis, where the ER probability increases (sharply too) with the mutation rate. We first seek the value$\mu_{*}$ at which ER occurs 50% of the time. This critical $\mu_{*}$, is the one above which ER becomes likely, so that the rescue probability is $P_{R}\left( \mu_{*} \right)=1/2$ and the ER rate is $\omega_{*}=-{\log\left( 1-P_{R}(\mu_{*}) \right)}/{N_{0}}={\log\left( 2 \right)}/{N_{0}}$. In this range of parameter ($\epsilon\ll1)$, we can use the approximate expressions in Eqs.(A9) and (A14) : $\omega\left( \mu_{*} \right)\approx2 \sqrt{{r_{max} \mu_{*}}/\pi}\exp(-r_{max} g\left( y_{D} \right)/\mu_{*})$. Solving for this equation yields a unique solution:

$$\begin{matrix} \mu_{*}\approx\frac{2 g\left( y_{D} \right) r_{\max}}{\mathcal{W}\left( \frac{8 g\left( y_{D} \right)}{\pi} \left( \frac{N_{0} r_{max}}{\log\left( 2 \right)} \right)^{2} \right)} \end{matrix} ,$$

(A15)

where $\mathcal{W}(.)$ is Lambert’s function. This rate can be approximated to a simpler form if we note that$N_{0}^{2}$ is typically very large compared to$g(y_{D})$ and$r_{max}$, which are of order 1. Therefore the denominator in Eq.(A15) is driven by the asymptotic limit of$\mathcal{W(.)}$ ($\mathcal{W}\left( v \right)\approx log(v/\log(v))$, for large$v$) and by the terms in$N_{0}$. Overall, to a reasonably good approximation:

$$\begin{matrix} \mu_{*}\underset{N_{0} \to\infty}{\approx} \frac{2 g\left( y_{D} \right) r_{\max}}{\log\left( \frac{N_{0}^{2}}{\log\left( N_{0}^{2} \right)} \right)} \end{matrix} .$$

(A16)

Recalling that$\mu=\sqrt{U \lambda}$, the critical mutation rate where the ER probability is 50% is

$$\begin{matrix} U_{*}\approx\frac{1}{\lambda}\left( \frac{2 g\left( y_{D} \right) r_{\max}}{\mathcal{W}\left( \frac{8 g\left( y_{D} \right)}{\pi} \left( \frac{N_{0} r_{max}}{\log\left( 2 \right)} \right)^{2} \right)} \right)^{2} \end{matrix}\underset{N_{0} \to\infty}{\approx}\frac{4 g\left( y_{D} \right)^{2} r_{\max}^{2}}{\lambda\log\left( \frac{N_{0}^{2}}{\log\left( N_{0}^{2} \right)} \right)^{2}} .$$

(A17)

This means that ER is only likely within a window of mutation rate $U_{*}\leq U\leq U_{max}$. This window narrows down as stress increases (increased decay rate and hence $g(y_{D})$, see **Supplementary Figure 6)** or as$N_{0}$ gets smaller, and its lower bound is roughly independent of dimensionality. The accuracy of this approximation is illustrated in **Figure 4** for the DN scenario ($g\left( y_{D} \right)=g_{DN}(y_{D})=\sqrt{y_{D}(1+y_{D})}-\cosh^{-1} \left( \sqrt{1+y_{D}} \right)$) and in **Supplementary Figure 5** below for the SV+DN scenario in the presence of standing variance ($g\left( y_{D} \right)=g_{DN+SV}(y_{D})=(y_{D}-\log\left( 1+y_{D} \right))/2$).

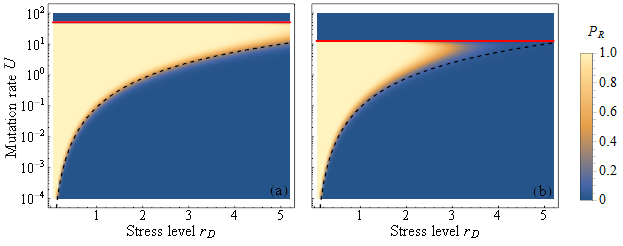

***Supplementary Figure 5:*** *same as* ***Figure 4*** *with standing variance plus de novo mutation.*

**VII. Height of the mutation window of ER**

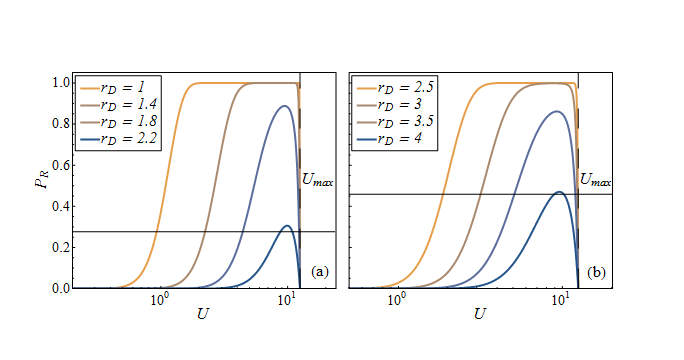

**Supplementary Figure 6:** Decrease of the width and height of the mutation window with stress. Colored plain lines show Eq.[6] for ER from *de novo* mutations (**a**) or from pre-existing standing genetic variance and *de novo* mutations (**b**). Black lines give $max(P_{R})$ from Eq.(A18) (for $r_{D}=2.2$ in (a) and $r_{D}=4$ in (b)). The colors and the parameters are the same as in **Figure 4** for increasing $r_{D}$.

Supplementary Figure 6 shows that, for sufficiently high stress$r_{D}$ or low inoculum size$N_{0}$, the ER probability cannot reach above some limited maximum$p= \max_{\mu\in\mathbb{R}^{+}} (P_{R})$. Figure 5 also shows that $max(P_{R})$ drops sharply (from $\max\left( P_{R} \right)=1$ to $\max\left( P_{R} \right)\ll1$) as$r_{D}$ increases or$N_{0}$ decreases. The goal of this section is to compute the values of$r_{D}$ where this transition occurs, namely the stress levels beyond which the population cannot avoid extinction, even if its mutation rate was to changed (e.g. by hyper-mutators).

The transition observed in Figure 5 occurs at some pairs of parameters $\{r_{D}^{*}(p),N_{0}^{*}(p)\}$ for which$\max\left( P_{R} \right)=p$ for some threshold value$p$. The shape of the transition in Figure 5 suggests that the values of $r_{D}^{*}(p)$ and $\log{(N}_{0}^{*}(p))$ are linearly related: so that$\log N_{0}^{*}(p)=a+b r_{D}^{*}(p)$ for some$(a,b)>0$. We also know that$P_{R}=1-\exp(-N_{0} \omega)$ where$\omega$ depends on $r_{D},r_{max},\mu,n$ (Eq.(5)). At $\{r_{D}^{*},\log N_{0}^{*}\}$ we thus have$p=\max\left( P_{R} \right)=1-\exp\left( -N_{0}^{*}(p) \omega^{*}(p) \right)$, where $\omega^{*}(p)$ is the corresponding ER rate along the transition of height$p$. This implies that$\log N_{0}^{*}(p)=\log(\left| \log\left( 1-p \right) \right|)-\log\omega_{*}\left( p \right)=a+b r_{D}^{*}(p)$, so that $\omega_{*}(p)=\omega\left( r_{D}^{*}\left( p \right) \right)=A exp(- B r_{D}^{*}\left( p \right))$ for some$(A,B)$. Since$\omega_{*}$ is independent of$N_{0}$ and is a maximum over$\mu$ (so independent of$\mu$), it must only depend on the remaining parameters$\{r_{D},r_{max},n\}$. The coefficients$A,B$, which determine its relationship with$r_{D}^{*}$, must thus only depend on $n$ and$r_{max}$. Therefore, we look for approximations of the form$\omega_{*}\approx A exp(- B r_{D}^{*})$, for some functions $A=A\left( n,r_{max} \right)$ and $B=B\left( n,r_{max} \right)$ for both $DN$ and $DN+SV$ scenarios. A simple fitting procedure, across a range of values of$\{n,r_{max}\}$, carried out with Matlab© Optimization Toolbox (the Matlab© code is available in supplementary file 2) suggests that $A\approx r_{max}/n$ and $B\approx\delta n/r_{max}$, leading to:

$$\begin{matrix} \max\left( P_{R} \right)=1-\exp\left( -N_{0}^{*}\left( p \right)\omega^{*}\left( p \right) \right) \\ \omega_{*}(p)=\frac{r_{max}}{n}\exp\left( -\delta\frac{n}{r_{max}} r_{D}^{*}(p) \right) \\ \mathrm{with}\left\{ \begin{matrix} DN :\delta=0.6 \\ DN+SV : \delta=0.31 \end{matrix} \right. \end{matrix}.$$

(A18)

From Eq.(A18) we can see that the height $max(P_{R})$ of the mutation window increases with an increase in the maximal growth rate $r_{max}$ or the population size $N_{0}$ and decreases with an increase in the dimensionality $n$ or the harshness of stress $r_{D}$ (the accuracy of this fitting for different values of $r_{max}$ and $n$ is illustrated in Supplementary figures 7). The figure below provides a comparison between the numerical value of $\max(P_{R})$ from Eq.[5] and the result of Eq.(A18).

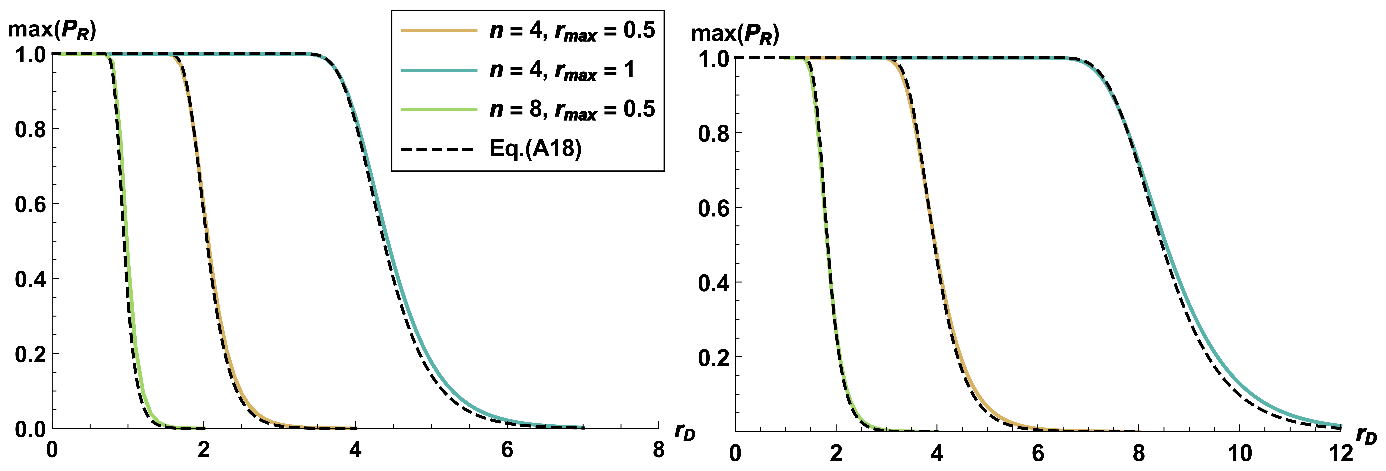

$$(a)$$

$$(b)$$

***Supplementary Figure 7:*** ${max(P}_{R})$ *as a function of*$r_{D}$ *for different values of the dimensionality and the maximal growth rate (given in legend), as computed from the ‘exact’ Eq.[5] vs. fitted approximation Eq.A(18). (a) Population adapting from de novo mutation only. (b) Population adapting from both de novo mutation standing genetic variance. In both cases* $N_{0}={10}^{5}$*.*

Solving $max(P_{R})=p=1-exp(-N_{0} \omega_{*}\left( p \right))$ in terms of$r_{D}^{*}(p)$ using Eq. A(18) yields the threshold stress at which$\max\left( P_{R} \right)=p$ :

$$r_{D}^{*}\left( p \right)=\frac{r_{max}}{n \delta}\log\left( \frac{N_{0} r_{max}}{n \left| \log\left( 1-p \right) \right|} \right)$$

$$.$$

(A19)

where$\delta$ is given in Eq. (A 18) depending on the scenario.

**VIII. Analytical and simulation models from Anciaux et al. (2018)**

*Explicit expression for a population initially clonal under the SSWM approximation from Anciaux et al. (2018)****:***

Here we provide the equations from Anciaux et al. (2018) for the rate of rescue per individual for a population initially clonal $\omega_{DN}$ in the SSWM regime ($U<U_{c}$). In this model, the distribution of fitness effects of mutations is also based on the FGM. Hence, from Eq.(2) of Anciaux et al. (2018), for a clonal population with a growth rate $-r_{D}$ in the new environment, the scaled growth rates of single-step mutants $y=r/r_{max}$, have the following probability density function:

$$f_{y}\left( y \right)=$$

$$\exp\left( - \frac{r_{max}\left( 2+y_{D}-y \right)}{\lambda} \right)\left( \frac{r_{max}}{\lambda} \right)^{n/2}\left( 1-y \right)^{n/2-1}\frac{{}_{0}F_{1}\left( \frac{n}{2} ,\left( \frac{r_{max}}{\lambda} \right)^{2}\left( 1+y_{D} \right)\left( 1-y \right) \right)}{\Gamma\left( n/2 \right)} ,$$

$$y\in]-\infty,1]$$

(A20)

with $y_{D}={r_{D}}/{r_{max}}\in[0,+\infty]$ and $r_{max}$, $n$, $\lambda$ defined in table 1. Where${}_{0}F_{1}(.,.)$ is the confluent hypergeometric function and $\Gamma\left( z \right)$ is the gamma function.

**F**ollowing Eq.(5) from Anciaux et al. (2018), the rate of rescue for a population initially clonal is:

$$\begin{matrix} \omega_{DN}=\frac{U}{r_{D}}\int_{0}^{1} \pi\left( y \right)f_{y}\left( y \right)dy \end{matrix} ,$$

(A21)

with $\pi\left( y \right)=1-exp(-2 y {r_{max}}/\sigma)$ the probability of establishment of a resistant genotype with scaled growth rate$y>0$ in the new environment (in figures 1 and 2 from the main text $\sigma=1$).

*Simulation algorithm from Anciaux et al. (2018)****:***

The following algorithm is described in the Methods section of Anciaux et al. (2018) and the explanations are directly extracted from the same section. The Mathematica code is provided in supplementary file I.

**while** COND

1. Mutation:

- Draw the number of mutations for each individual from a Poisson distribution with rate $U$
- add mutation effects (when relevant) to the current parent phenotype, by draowing these into a multivariate normal distribution $N\left( \boldsymbol{0}, \lambda\mathbf{I}_{\boldsymbol{n}} \right)$

2. Selection and drift:

- Genotypes forming the new generation are sampled with replacement from the previous one with weight $W_{i}=e^{r_{i}}$ for the genotype $i$.

3. Demographic stochasticity:

- The size $N_{t+1}$ of population at generation$t+1$ is drawn as a Poisson number $N_{t+1} \sim Poisson(N_{t} \bar{W})$, with$\bar{W}=\bar{exp(r)}$ the mean Darwinian fitness ($W=exp(r)$) and$N_{t}$ the population size, in the previous generation.

**end while**

With COND = **if** ($\bar{r}_{t}>0$, $\exp\left( -2 N_{t} \bar{r}_{t} \right)<{10}^{-12}$) & $N_{t}>0$. A population is considered rescued when it reaches a population size$N_{t}$ and mean growth rate$\bar{r}_{t}$ such that its ultimate extinction probability, if it were monomorphic and stable over time (probability $\exp\left( -2 N_{t} \bar{r}_{t} \right)$), would lie bellow${10}^{-12}$. This is a conservative criterion: once$\bar{r}_{t}$ has become positive, we expect it to remain so, yielding further increases in population size and thus further decreasing the probability of future extinction.

Points 2 and 3 from the algorithm correspond to reproduction and have been separated into 2 separate phases to increase the speed of the algorithm. However, this is exactly equivalent to drawing independent Poisson reproductive outputs from each individual.

In the algorithm in supplementary file 1, the effects of mutations are not drawn every generations as in point 1. Instead a large number (${10}^{6}$) of mutation effects are drawn before the simulations and mutations effects are drawn in this “pre-set” every generations to increase the speed of the algorithm.

For rescue from populations at mutation-selection balance, $10$ replicate initial equilibrium populations were generated, each by starting from an optimal clone and running the same algorithm with fixed population size ($N_{t}={10}^{6}$) until the mean growth rate had visually stabilized to a fixed value (close to its theoretical equilibrium value$\bar{r}{}_{eq}=r_{max}-U$ (for$U<U_{c}$) for more than $1000$ generations. Then the optimum was shifted by$\sqrt{2(r_{D}+r_{max})}$ phenotypic units, and $1000$ replicate ER simulations were performed (same algorithm as for de novo rescue), from each of the $8$ replicate equilibrium populations.

**Supplementary figure 8** shows the dynamic of the mean fitness and the population size of $4$ replicates ($2$ rescues and $2$ extinctions) of a simulated population using the above algorithm for a population with high mutation rate ($U>U_{c}$).

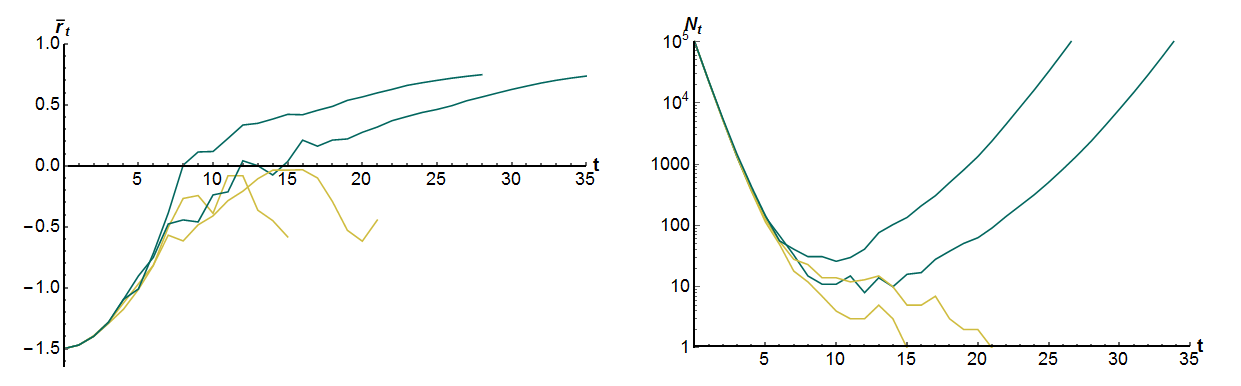

A

B

**Supplementary Figure 8:** Dynamics of the mean fitness$\bar{r}_{t}$ (A) and the population size$N_{t}$ (B) of replicate populations starting from a clone at $-r_{D}=-1.5$ at size $N_{0}={10}^{5}$. The blue lines represents rescues and the yellow lines extinctions. Parameters for the simulations are $r_{max}=1$, $U=100 U_{c},$ $n=4$ and $\lambda=5\times{10}^{-3}$.

*.*
