## Appendix II for "Population persistence under high mutation rate: from evolutionary rescue to lethal mutagenesis"

### Appendix II : Time-inhomogeneous Feller diffusion

#### 1 Time-inhomogeneous diffusions

Let $(X_t; t \geq 0)$ be a time-inhomogeneous diffusion process which is the unique solution to the following SDE

$$dX_t = a(t; X_t)\, dt + \sqrt{b(t; X_t)}\, dB_t,$$

where $b(t, x) > 0$ for all $t, x$, and both functions $a$ and $b$ are continuous in $t$ and Lipschitz-continuous in $x$. Now let $g : \mathbb{R} \to \mathbb{R}$ be any twice differentiable function with bounded derivatives and set

$$p_s(t; x) := E(g(X_s)|X_t = x) \qquad s > t.$$

Let $L_t$ denote the generator of $X$ at time $t$, that is, for any twice differentiable $f : \mathbb{R} \to \mathbb{R}$ with locally bounded derivatives,

$$L_t f(x) := \lim_{\varepsilon \uparrow 0} \varepsilon^{-1} E(f(X_{t+\varepsilon}) - f(X_t)|X_t = x) = a(t; x) f'(x) + \frac{1}{2} b(t; x) f''(x).$$

Then it is well-known (Bansaye and Simatos, 2015) that $p_s$ is differentiable in $t$ and twice differentiable in $x$, and satisfies

$$-\frac{\partial p_s}{\partial t}(t; x) = L_t p_s(t, x),$$

where it is implicit that in the right-hand side, the generator applies to the second variable of $p_s$ and the first one is fixed equal to $t$, that is,

$$-\frac{\partial p_s}{\partial t}(t; x) = a(t; x)\frac{\partial p_s}{\partial x}(t; x) + \frac{1}{2} b(t; x)\frac{\partial^2 p_s}{\partial x^2}(t; x). \tag{1}$$

#### 2 The case of inhomogeneous Feller diffusions

Here we assume that $a(t; x) = r(t)x$ and $b(t; x) = \sigma(t)x$, so that $X$ is a *Feller diffusion* with time-dependent growth rate $r(t)$ and variance $\sigma(t)$, both assumed continuous in $t$. Note that 0 is an absorbing point, which is accessible as soon as $\sigma$ is strictly positive, which we assume. Now we apply the results of the previous section to $g(x) := e^{-\lambda x}$.

**Theorem 1.** *For any $t < s$, we have*

$$E\left(e^{-\lambda X_s} | X_t = x\right) = e^{-x\varphi_{\lambda,s}(t)} \qquad x \geq 0,$$

*where*

$$\varphi_{\lambda,s}(t) = \left(\frac{1}{\lambda} e^{-\int_t^s r(u)\,du} + \frac{1}{2}\int_t^s \sigma(u) e^{-\int_t^u r(v)\,dv} du\right)^{-1}.$$

*Proof of the Theorem.* It is well-known that $X$ satisfies the branching property, in the sense that the sum of two independent copies of $X$ started respectively at $x$ and $y$, has the same law as $X$ started at $x + y$. This ensures that for any $s > t \geq 0$ and $\lambda > 0$, there exists a non-negative real number $\varphi_{\lambda,s}(t)$ such that

$$p_s(t; x) = e^{-x\varphi_{\lambda,s}(t)} \qquad x \geq 0.$$

Thanks to (1), $\varphi_{\lambda,s}$ is differentiable at any $t < s$ and we have the following equalities on $[0, s)$

$$-\frac{\partial p_s}{\partial t}(t; x) = x\varphi'_{\lambda,s}(t) e^{-x\varphi_{\lambda,s}(t)},$$

$$\frac{\partial p_s}{\partial x}(t; x) = -\varphi_{\lambda,s}(t) e^{-x\varphi_{\lambda,s}(t)}$$

and

$$\frac{\partial^2 p_s}{\partial x^2}(t; x) = (\varphi_{\lambda,s}(t))^2 e^{-x\varphi_{\lambda,s}(t)}$$

Then thanks to (1) again, we get

$$\varphi'_{\lambda,s} = -r\varphi_{\lambda,s} + \frac{\sigma}{2}\varphi^2_{\lambda,s} \tag{2}$$

We are now going to solve (2) with the terminal condition $\lim_{t\uparrow s} p_s(t; x) = e^{-\lambda x}$, that is,

$$\lim_{t\uparrow s} \varphi_{\lambda,s}(t) = \lambda$$

Assume that $\varphi_{\lambda,s}$ satisfies (2). Set

$$R_s(t) := \int_t^s r(u)\,du \qquad t < s$$

and

$$\psi_{\lambda,s}(t) := \varphi_{\lambda,s}(t) e^{-R_s(t)} \qquad t < s.$$

Then we have $R'_s = -r$ and

$$\psi'_{\lambda,s} = \varphi'_{\lambda,s} e^{-R_s} + r\varphi_{\lambda,s} e^{-R_s} = \frac{\sigma}{2}\varphi^2_{\lambda,s} e^{-R_s} = \frac{\sigma}{2}\psi^2_{\lambda,s} e^{R_s}.$$

Now we integrate $\psi'_{\lambda,s} = \frac{\sigma}{2}\psi^2_{\lambda,s} e^{R_s}$ in the following way. Let us assume that there is $t_0 < s$ such that $\psi_{\lambda,s}(t_0) = 0$. Then $\psi'_{\lambda,s}(t_0) = 0$ so by uniqueness of the solution, $\psi_{\lambda,s}$ is zero on $[t_0, s)$. But this would contradict the fact that $\lim_{t\uparrow s} \psi_{\lambda,s}(t) = \lim_{t\uparrow s} \varphi_{\lambda,s}(t) = \lambda > 0$. So $\psi_{\lambda,s}(t) > 0$ for any $t < s$ and we can write

$$\int_t^s \frac{\psi'_{\lambda,s}(u)}{\psi_{\lambda,s}(u)^2}\, du = \frac{1}{2}\int_t^s \sigma(u) e^{R_s(u)} du,$$

where the left-hand side can be integrated into

$$\left[-\frac{1}{\psi_{\lambda,s}}\right]_t^s = -\frac{1}{\lambda} + \frac{1}{\psi_{\lambda,s}(t)},$$

due to the assumption that $\lim_{u\uparrow s} \psi_{\lambda,s}(u) = \lambda$. So we finally get

$$\begin{aligned}\frac{1}{\varphi_{\lambda,s}(t)} = \frac{e^{-R_s(t)}}{\psi_{\lambda,s}(t)} = \frac{e^{-R_s(t)}}{\lambda} + \frac{1}{2}\int_t^s \sigma(u) e^{R_s(u)-R_s(t)} du \\ = \frac{1}{\lambda} e^{-\int_t^s r(u)\,du} + \frac{1}{2}\int_t^s \sigma(u) e^{-\int_t^u r(v)\,dv} du,\end{aligned}$$

which yields the expected result. □

From now on, we fix $t = 0$ and write $\varphi_\lambda(s) := \varphi_{\lambda,s}(0)$ and $\varphi(s) := \varphi_{\infty,s}(0)$, that is

$$\varphi_\lambda(s) = \left(\frac{1}{\lambda} e^{-\int_0^s r(u)\,du} + \frac{1}{2}\int_0^s \sigma(u) e^{-\int_0^u r(v)\,dv} du\right)^{-1}$$

and

$$\varphi(s) = \left(\frac{1}{2}\int_0^s \sigma(u) e^{-\int_0^u r(v)\,dv} du\right)^{-1}.$$

**Corollary 2.** *Starting at $X_0 = x$, the state $X_s$ of the inhomogeneous Feller diffusion can be written as*

$$X_s = \sum_{i=1}^{N_s} Y_i(s),$$

*where $N_s$ is a Poisson random variable with parameter $x\varphi(s)$ and the $Y_i(s)$ are i.i.d. exponential random variables independent of $N_s$, with common parameter $\rho(s)$, where*

$$\rho(s) := \left(\frac{1}{2}\int_0^s \sigma(u) e^{\int_u^s r(v)\,dv} du\right)^{-1}.$$

*In particular, if $T$ denotes the hitting time of 0, also called* extinction time, *then*

$$\mathbb{P}(T < s | X_0 = x) = e^{-x\varphi(s)}.$$

*Proof.* The last part of the corollary just stems from the following argument

$$\mathbb{P}(T < s | X_0 = x) = \mathbb{P}(X_s = 0 | X_0 = x) = \lim_{\lambda\to\infty} E\left(e^{-\lambda X_s} | X_0 = x\right) = e^{-x\varphi(s)}.$$

Now set

$$Z_s := \sum_{i=1}^{N_s} Y_i(s)$$

Let us prove that $Z_s$ has the same Laplace transform as $X_s$, that is

$$\mathbb{E}\left(e^{-\lambda Z_s}\right) = e^{-x\varphi_\lambda(s)} = \mathbb{E}\left(e^{-\lambda X_s}\right),$$

which will prove the first part of the corollary. It is elementary that

$$\begin{aligned}\mathbb{E}\left(e^{-\lambda Z_s}\right) &= \mathbb{E}\left[\mathbb{E}\left(e^{-\lambda Y_1}\right)^{N_s}\right] \\ &= \exp\left(-x\varphi(s)\left[1 - \mathbb{E}\left(e^{-\lambda Y_1}\right)\right]\right) \\ &= \exp\left(-x\varphi(s)\left[1 - \frac{\rho(s)}{\lambda + \rho(s)}\right]\right)\end{aligned}$$

Now it is easy to see that

$$\varphi(s)\left[1 - \frac{\rho(s)}{\lambda+\rho(s)}\right] = \frac{\lambda\,\varphi(s)}{\lambda+\rho(s)} = \frac{\varphi(s)/\rho(s)}{\frac{1}{\rho(s)} + \frac{1}{\lambda}} = \frac{e^{\int_0^s r(u)\,du}}{\frac{1}{2}\int_0^s \sigma(u) e^{\int_u^s r(v)\,dv} du + \frac{1}{\lambda}} = \varphi_\lambda(s),$$

which ends the proof. □

### 3 Large time convergence

Now we assume that

$$\int_0^\infty \sigma(u)\, e^{-\int_0^u r(v)\,dv} du < \infty,$$

so that $\varphi(\infty) > 0$. Recall that $Y_1(s)$ is exponential with expectation $1/\rho(s)$, so that

$$Y_1(s)\, e^{-\int_0^s r(u)\,du}$$

is exponential with expectation

$$e^{-\int_0^s r(u)\,du}/\rho(s) = \frac{1}{2}\int_0^s \sigma(u)\, e^{-\int_0^u r(v)\,dv} du,$$

which by assumption converges to $1/\varphi(\infty) < \infty$. This can be recorded in the following statement, where we define $a := \varphi(\infty)$.

**Proposition 3.** *Set*

$$a := \left( \frac{1}{2} \int_0^\infty \sigma(u)\, e^{-\int_0^u r(v)\,dv} du \right)^{-1} > 0.$$

*Then conditional on* $X_0 = x$*,* $X_s\, e^{-\int_0^s r(u)\,du}$ *converges in distribution to the terminal random variable*

$$W := \sum_{i=1}^{N} Y_i,$$

*where* $N$ *is a Poisson random variable with parameter* $ax$ *and the* $Y_i$ *are i.i.d. exponential random variables independent of* $N$*, with common parameter* $a$*.*

Notice that the terminal random variable $W$ is 0 iff $N = 0$, which occurs with probability $e^{-ax}$. Since this is also the probability that $T < \infty$ and that $\{T < \infty\} \subset \{N = 0\}$, these two events actually coincide with probability 1.

Also note that actually the convergence in the previous proposition holds pathwise (i.e. with probability 1, or 'for almost all realizations'). This is due to the fact that $(X_s\, e^{-\int_0^s r(u)\,du}; s \geq 0)$ is actually a martingale.
