## Supplementary material for "Population persistence under high mutation rate: from evolutionary rescue to lethal mutagenesis": Matlab code for curve fitting: curve_fit.m

```
clear all

%%% curve fitting: finds the coefficients in the formula of max PR

%%% case 1: no standing variation

% functions: general form
fomega=@(a,b,rmax,n,rD)a*rmax/n*exp(-n/rmax*b*rD);
fPR = @(a,b,rmax,n,rD,N0)(1-exp(-N0*fomega(a,b,rmax,n,rD)));

% loads the numerical values of max PR (numerical values obtained with the script num_comp_maxPR.m)
rmax=0.5; n=4;
load(['VPR_rmax_' num2str(rmax) '_n_' num2str(n) '.mat'],'VrD','VPR');
VPR1=VPR;
VrD1=VrD;
rmax=1; n=4;
load(['VPR_rmax_' num2str(rmax) '_n_' num2str(n) '.mat'],'VrD','VPR');
VPR2=VPR;
VrD2=VrD;
rmax=0.5; n=8;
load(['VPR_rmax_' num2str(rmax) '_n_' num2str(n) '.mat'],'VrD','VPR');
VPR3=VPR;
VrD3=VrD;

% functional to be minimized
f=@(A)norm(fPR(A(1),A(2),0.5,4,VrD1,1e5)-VPR1)^2+norm(fPR(A(1),A(2),1,4,VrD2,1e5)-VPR2)^2+norm(fPR(A(1),A(2),0.5,8,VrD3,1e5)-VPR3)^2;
options = optimoptions('fmincon','Display','off');
xnosv = fmincon(f,[0 0],[],[],[],[],[1 0],[10 10],[],options); % minization. 
display('parameter values, no standing variation')
xnosv

%%% case 2: with standing variation

% loads the numerical values of max PR (numerical values obtained with the script num_comp_maxPRSV.m)
rmax=0.5; n=4;
load(['VPRSV_rmax_' num2str(rmax) '_n_' num2str(n) '.mat'],'VrD','VPR');
VPR1=VPR;
VrD1=VrD;
rmax=1; n=4;
load(['VPRSV_rmax_' num2str(rmax) '_n_' num2str(n) '.mat'],'VrD','VPR');
VPR2=VPR;
VrD2=VrD;
rmax=0.5; n=8;
load(['VPRSV_rmax_' num2str(rmax) '_n_' num2str(n) '.mat'],'VrD','VPR');
VPR3=VPR;
VrD3=VrD;

% functional to be minimized
f=@(A)norm(fPR(A(1),A(2),0.5,4,VrD1,1e5)-VPR1)^2+norm(fPR(A(1),A(2),1,4,VrD2,1e5)-VPR2)^2+norm(fPR(A(1),A(2),0.5,8,VrD3,1e5)-VPR3)^2;
xsv = fmincon(f,[0 0],[],[],[],[],[1 0],[10 10],[],options); % minization. 
display('parameter values, no standing variation')
xsv
```
