## Supplementary material for "Population persistence under high mutation rate: from evolutionary rescue to lethal mutagenesis": Matlab code for curve fitting: num_comp_maxPRSV.m

```
function calcul_matPRSV(rmax,n)

%%% numerical computation of the values of mu that maximize PR, and
%%% saves the corresponding values of PR et omega
%%% case with standing variation

nom=['VPRSV_rmax_' num2str(rmax) '_n_' num2str(n) '.mat' ];
i=0; 
for rD=0.1:0.01:15
    i=i+1
    VrD(i)=rD;
    [mumax omega PR]=maxPRSV(rmax,n,rD,1e5);
    Vmumax(i)=mumax;
    Vomega(i)=omega;
    VPR(i)=PR;
end
save(nom)

function [mumax omega PR]=maxPRSV(rmax,n,rD,N0)

%%% based on the numerical computation of max omega

yD=rD/rmax;
f=@(y)y-(1+yD).*(1-exp(-2*y))/2;
h=@(y)exp(y);

Den=@(y,mu)h(y).^(n/2).*exp(-rmax./mu.*f(y));
H=@(y,mu)h(y).^(n/2).*exp(-rmax./mu.*f(y)).*(mu/rmax-f(y));
as=1e-3;
dx=1e-2; X=0:dx:100;
H0=sum(H(X,as));
while H0>0
    as=as+1e-3;
    H0=sum(H(X,as));
end
% max omega is reached when H changes sign

mumax=as;
omega=2*as/(sum(Den(X,as))*dx);
PR=1-exp(-N0*omega);
```
