## Supplementary material for "Population persistence under high mutation rate: from evolutionary rescue to lethal mutagenesis": Matlab code for curve fitting: plot_curves_noSV.m

```
%%% plot the curves corresponding to maxPR vs rD
%%% analytical formula vs numerical values
%%% case without standing variation

close all
clear all

% analytical formula
alpha=1; beta=0.59;
omegas=@(rmax,n,rD)alpha*rmax/n*exp(-n/rmax*beta*rD);
valmaxPR=@(rmax,n,rD,N0)1-exp(-N0*omegas(rmax,n,rD));

%%%

rmax=1/2;n=4;
nom=['VPR_rmax_' num2str(rmax) '_n_' num2str(n) '.mat' ];
load(nom)
N0=1e5;
j=0;
for rD=VrD
    j=j+1;
    s(j)=valmaxPR(rmax,n,rD,N0);
end
hold on
h1=plot(VrD,VPR,'b-','LineWidth',1.5);
plot(VrD,s,'b--','LineWidth',1.5);

%%%

rmax=1/2;n=8;
nom=['VPR_rmax_' num2str(rmax) '_n_' num2str(n) '.mat' ];
load(nom)
N0=1e5;
j=0;
for rD=VrD
    j=j+1;
    s(j)=valmaxPR(rmax,n,rD,N0);
end
h2=plot(VrD,VPR,'g','LineWidth',1.5);
plot(VrD,s,'g--','LineWidth',1.5)

%%%

rmax=1;n=4;
nom=['VPR_rmax_' num2str(rmax) '_n_' num2str(n) '.mat' ];
load(nom)
N0=1e5;
j=0;
for rD=VrD
    j=j+1;
    s(j)=valmaxPR(rmax,n,rD,N0);
end
h3=plot(VrD,VPR,'r','LineWidth',1.5);
plot(VrD,s,'r--','LineWidth',1.5)

legend([h1,h3,h2],'n=4, r_{max}=1/2','n=4, r_{max}=1','n=8, r_{max}=1/2')
set(gca,'FontSize',18,'FontName','Times New Roman')
xlabel('r_D','FontAngle','italic')
ylabel('max PR','Interpreter','Latex')
```
