## Supplementary material for "Population persistence under high mutation rate: from evolutionary rescue to lethal mutagenesis": Matlab code for curve fitting: readme.txt

%%% matlab codes:
num_comp_maxPR.m   % numerical computation of the values of mu that maximize PR (no standing variation)
num_comp_maxPRSV.m  %  numerical computation of the values of mu that maximize PR (with standing variation)    
curve_fit.m % curve fitting of max PR          
plot_curves_SV.m  % graphical comparison between analytical formula and numerical values (with standing variation)
plot_curves_noSV.m  % plot_curves_SV.m  % graphical comparison between analytical formula and numerical values (no standing variation)
  
%%% other files: numerical values of max PR obtained with num_comp_maxPR.m and num_comp_maxPRSV.m 
VPR_rmax_0.5_n_4.mat
VPR_rmax_1_n_4.mat
VPR_rmax_0.5_n_8.mat
VPRSV_rmax_0.5_n_4.mat
VPRSV_rmax_0.5_n_8.mat
VPRSV_rmax_1_n_4.mat
